# Solvent-free Nanoparticle Assembly Protocol (SNAP): one-pot formulation of drug loaded polyester nanoparticles and their vessel size-dependent perivascular transport in the brain

**DOI:** 10.64898/2026.06.29.735299

**Authors:** Elena A. Andreyko, Miles Pourbaghi, Sarah E. Stabenfeldt, Rachael W. Sirianni

**Author notes:** **Author to whom correspondence should be addressed:** Rachael Sirianni, PhD, Department of Biochemistry and Molecular Biotechnology, UMass Chan Medical School, 364 Plantation Street, Worcester, MA 01605.

## Abstract

This work describes a new approach for rapid and reproducible formulation of drug loaded biodegradable nanoparticles based on polyester copolymers, including poly(lactic acid)-poly(ethylene glycol) (PLA-PEG) and poly(caprolactone)-poly(ethylene glycol) (PCL-PEG). The new approach, termed Solvent-free Nanoparticle Assembly Protocol (SNAP), carries several advantages over conventional polyester formulation strategies, including very rapid formulation (minutes) and the ability to use nanoparticles immediately without lengthy solvent evaporation or washing steps. Altering polyester molecular weight and concentration, alongside the introduction of specific functional groups yielded precise control of nanoparticle properties, including size, shape, surface charge, drug release and loading. We examined loading of multiple therapeutic compounds, including diclofenac, loperamide, bortezomib, CT179, panobinostat, docetaxel, methotrexate, and camptothecin. The SNAP protocol facilitated the rapid production of stable, drug-loaded nanoparticles with a narrow size distribution and generally good drug loading. Using Fluorescence Resonance Energy Transfer (FRET) and size exclusion chromatography (SEC) with a focus on the model agent Rhodamine B, we were able to carefully examine stability of the nanoparticle and assess the distribution of small molecules within the polymer as well as nanoparticle stability. *In vivo* evaluation of fluorescently labeled nanoparticles using real-time, intravital microscopy showed that, after direct administration to cerebrospinal fluid (CSF) via the intrathecal cisterna magna (IT-CM) route, the dynamic accumulation of nanoparticles within the perivascular space (PVS) depends on the size of the vessel that is imaged. Nanoparticles accumulated steadily within the PVS of large vessels, while accumulating more slowly and exhibiting clearance from medium-sized and smaller vessels over the course of several hours. In sum, these studies present a new platform for facile production of polyester nanoparticles, demonstrate their ability to encapsulate a variety of hydrophobic small molecules, and expand our knowledge on the development of nanocarriers for intrathecal administration. Taken together, these data open new opportunities for development safer and more effective nanoparticle-based therapies.

## Introduction

Nanoparticles (NPs) are synthetic or natural colloids between 1 to 500 nm in size, and they can offer many advantages for drug delivery due to their ability to protect bioactive agents from degradation or clearance within physiological environments [1, 2]. Encapsulation of drug within a nanoparticle can offer several advantages, including reduced toxicity, the ability to target specific cells and tissues, and enhanced efficacy [3, 4, 5]. Although some polymeric nanoparticle formulations have found clinical success [6, 7], several obstacles have prevented widespread adaptation of polymeric systems as drug carriers. The streamlined production of polymeric NPs is impeded by critical manufacturing obstacles. Batch-to-batch variability remains a primary concern, as maintaining uniform particle size, polydispersity, surface charge, and morphology is inherently difficult. Furthermore, the physical and chemical stability of both the polymer and sensitive payloads such as proteins or RNA is often compromised by high-energy fabrication methods like sonication or homogenization. Finally, current methodologies are burdened by process-intensive requirements, including the hazardous use of organic solvents necessitating additional, time-consuming purification steps like evaporation, dialysis, or lyophilization.

In this study, we developed a new, Solvent-free Nanoparticle Assembly Protocol (SNAP) for one-pot rapid, solvent-free, reproducible, and customizable formulation of drug loaded polyester NPs. We hypothesized that altering key formulation parameters, including molecular weight, concentration, nature of polymers, as well as biophysical features of the encapsulated payload, would lead to the production of NPs with defined, controllable properties (size, shape, surface charge, drug release and loading). To test this hypothesis, we created over twenty new drug loaded NPs with different properties. By optimizing di-block copolymer concentration and molecular weight, we developed small spherical nanocarriers. We showed that for effective encapsulation, the polymer’s core must be functionalized with additional functional groups for payload interaction, while the drug itself should be small and hydrophobic. Intravital microscopy demonstrated that these particles exhibit extended circulation and a gradual accumulation/clearance vessel size-dependent pattern in the perivascular regions of the brain. The new SNAP technique allows for quick, customizable synthesis of drug-loaded NPs, offering enhanced opportunities for therapeutic delivery.

## Results

### 1. Preparation and characterization of Drug empty SNAP NPs

The Solvent-free Nanoparticle Assembly Protocol (SNAP) was developed to prepare NPs based on the block copolymers poly(ethylene glycol)-b-poly(D,L-lactide) (PLA-PEG) and poly(ethylene glycol)-b-poly(caprolactone) (PCL-PEG). PLA and PCL are hydrophobic polyesters that are widely used to generate biodegradable and biocompatible NPs. The use of amphiphilic polyesters enabled us to generate a stable, hydrophobic nanoparticle core coated with a hydrophilic PEG shell; this formulation scheme is illustrated in **Fig. 1A**. To generate NPs via self-assembly, amphiphilic copolymers were dissolved in diethylene glycol monoethyl ether, and cold water was added dropwise under stirring (**Fig. 1B**). Diethylene glycol monoethyl ether plays a unique role in this formulation, being both a solvent that can solubilize hydrophobic polymers, but also being a biocompatible component of the formulation that does not need to be removed prior to use *in vivo* [8, 9]. The SNAP approach reduces formulation time by 97% compared to traditional methods by facilitating rapid NP formation within 5 minutes of mixing, eliminating the need for post-synthetic solvent evaporation or thermal treatment.

**Figure 1.**
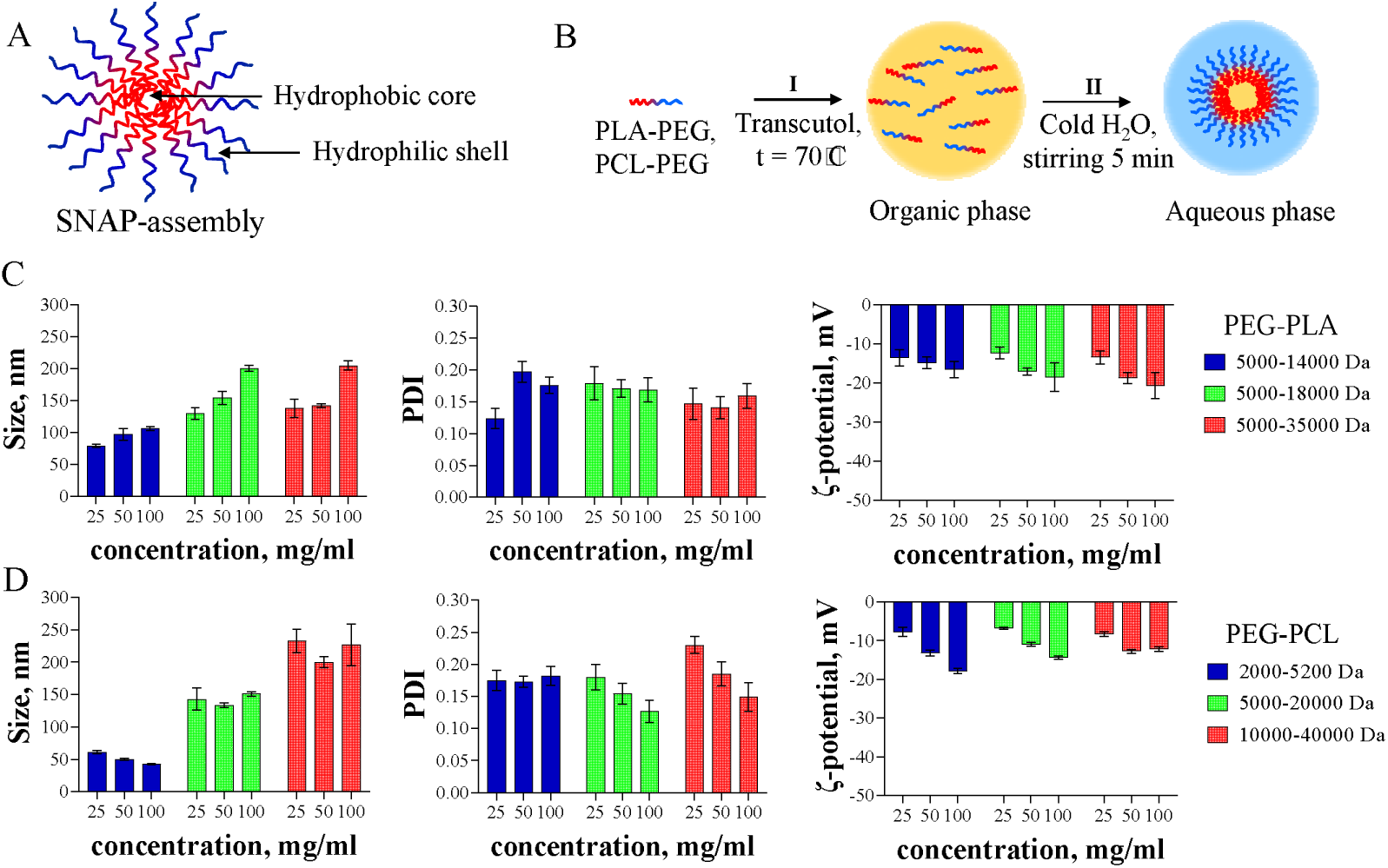
Schematic representation of (A) SNAP NPs, (B) drug-empty NPs formulation. Plot of (C) mean size, PDI, ζ-potential for drug-empty NPs formed by PLA-PEG. Plot of (D) mean size, PDI, ζ-potential for drug-empty NPs formed by PCL-PEG.

To study how formulation conditions influence the properties of resultant NPs, SNAP was used to formulate NPs while varying the type of constituent polymers, their molecular weight, and their concentration in diethylene glycol monoethyl ether. We observed that each of these parameters can be used to adjust the hydrodynamic diameter of resulting NPs. For example, increasing the molecular weight of PLA-PEG from 14kDa-5kDa to 35kDa-5kDa increasing polymer concentration from 25 to 100 mg/mL in diethylene glycol monoethyl ether yielded an increase in diameter of resultant NPs size from 79.2±2.3 to 204.5±7.2 nm (**Fig. 1C**). However, for PCL-PEG NPs only changing molecular weight of polymer led to an increase in the NPs size (**Fig. 1E**). The low value (<0.2) of polydispersity index (PDI) for all formulations indicates that all systems have narrow size distributions (**Fig. 1C and D**). Negative value of surface charge from −6.8±0.3 to −20.7±3.3 mV measured by dynamic light scattering showed that all NPs are stable (**Fig. 1C and D**). The lowest absolute value of ζ-potential from −6.8±0.3 to −13.6±2.1 mV was observed when polymer concentration was low (25mg/mL in diethylene glycol monoethyl ether) for all types of polymers with different molecular weights (**Fig. 1C and D**).

### 2. Loperamide Loaded SNAP NPs

To assess the ability of NPs formulated via SNAP to load drugs, we focused on loperamide, which is a small molecule that is commonly used as a model compound in CNS drug delivery work. Loperamide is a potent lipophilic opioid, however, it does not cross the blood-brain barrier (BBB) at normal therapeutic doses. Additionally, loperamide is poorly soluble in water (0.0014 g/mL at pH 7.1), which limits deliverable dose. To generate drug loaded NPs with the SNAP approach, amphiphilic copolymers (PLA-PEG or PCL-PEG) at concentrations 25, 50 and 75 mg/ml were dissolved in diethylene glycol monoethyl ether and mixed with 1 mg of loperamide followed by nanoprecipitation into the cold water (**Fig. 2A**). In this study, loperamide loaded SNAP NPs were formulated using two types of polymers with different molecular weights and concentration in diethylene glycol monoethyl ether to investigate whether these parameters will impact on NPs properties (size, ζ-potential, PDI) and drug loading. In contrast to what was observed for drug-empty NPs, we observed minimal to no effect of polymer concentration on the particle size, ζ-potential and PDI (**Fig. 2B and C**). Increasing the molecular weight of PLA-PEG from 5kDa-14kDa to 5kDa-34kDa and PCL-PEG from 2kDa-5.2kDa to 10kDa-40kDa increased NP size from 43.8±1.5nm to 190.8±6.5 nm (**Fig. 2B and C**). Results indicated that there were no significant changes in loperamide loading as a function of molecular weight (**Fig. 2B and C**). Drug loading was generally modest (<0.5 w/w%), with the highest loading (1.2±0.4 w/w%) observed for NPs composed of PEG-PLA (5kDa-14kDa) and formulated at the lowest concentration 25 mg/ml in diethylene glycol monoethyl ether (**Fig. 2B**). To increase the loperamide loading, we also generated NPs using PEG-PLA-COOH, which bears carboxylic groups that can participate in Lewis acid (PEG-PLA-COOH) - base (loperamide) interaction. In these conditions, we were able to increase loperamide loading to 2.1±0.2 w/w% (**Fig. 2B**). The Loperamide loaded PEG-PLA-COOH NPs were 90.8±12.1 nm in size and spherical, as characterized by dynamic light scattering (DLS) (**Fig. 2B**) and transmission electron microscopy (TEM) (**Fig. 4B**). The absolute value of ζ-potential at pH=7 is higher for loperamide loaded NPs based on PLA-PEG compared to PCL-PEG and drug empty NPs, indicating their increased stability in solution.

**Figure 2.**
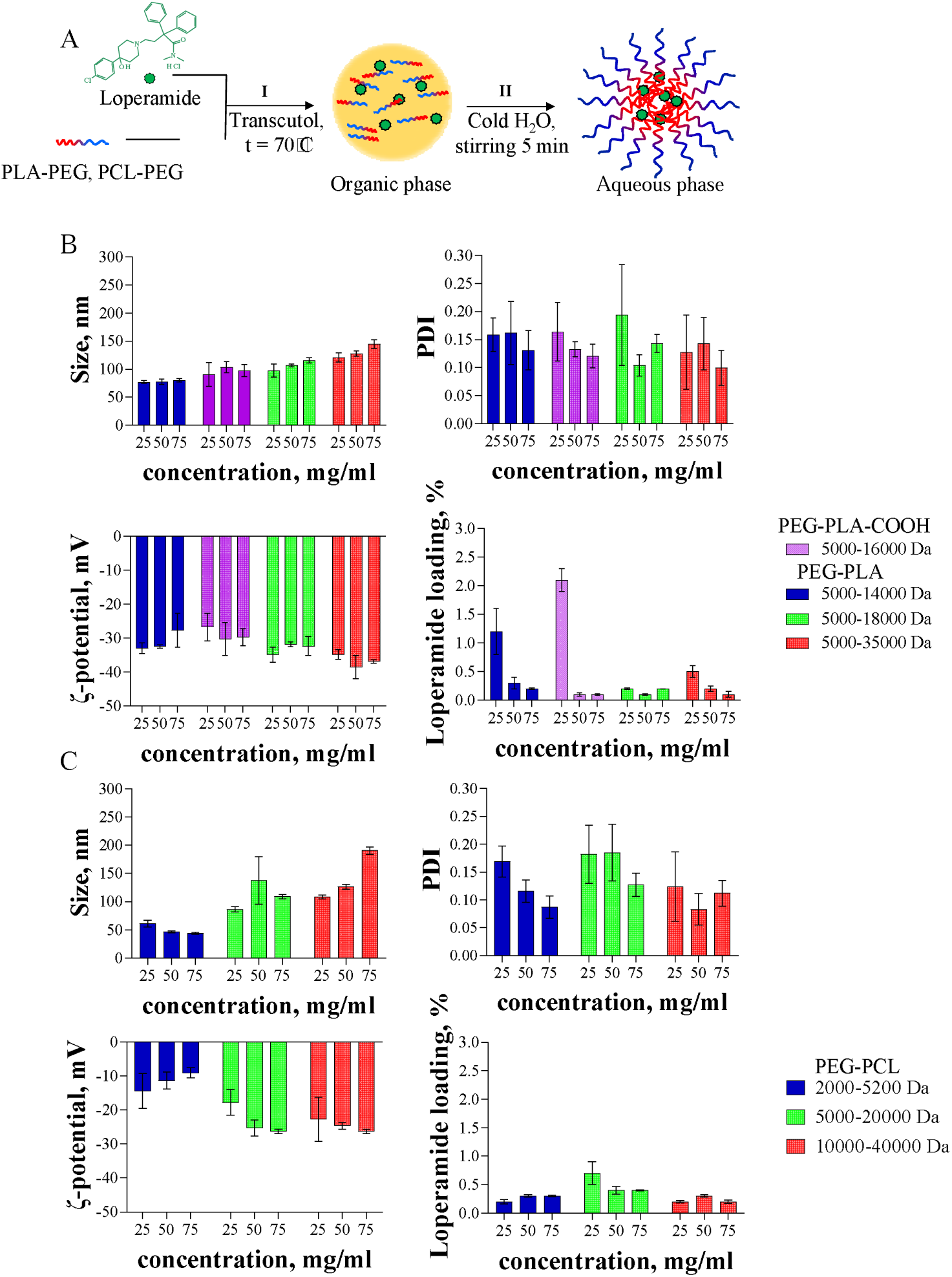
(A) Schematic representation of SNAP NPs formulation with drugs. Plot of (B) mean size, PDI, ζ-potential and loperamide loading for NPs formed by PLA-PEG. Plot of (C) mean size, PDI, ζ-potential and loperamide loading for NPs formed by PCL-PEG.

Drug that is solubilized after formulation with a NP system can be contained within the NP core or bound to its surface, or drug may exist freely solubilized in solution; this is a particular concern for loperamide, which is a molecular that is both lipophilic and partially soluble in water. We therefore developed approaches to determine the quantity of loperamide that is freely solubilized within the formulation with size exclusion high-performance liquid chromatography (SE-HPLC). Drug-empty NPs and NPs loaded with loperamide that were not washed served as negative and positive controls, respectively. We observed that the loperamide detected in unwashed NPs primarily exists in freely solubilized form (**Fig. 3A and C**). Once NPs were washed three times, unencapsulated loperamide was removed from the system, leaving 7.0±1.5% of the drug effectively entrapped within the system. (**Fig. 3B and D**).

**Figure 3.**
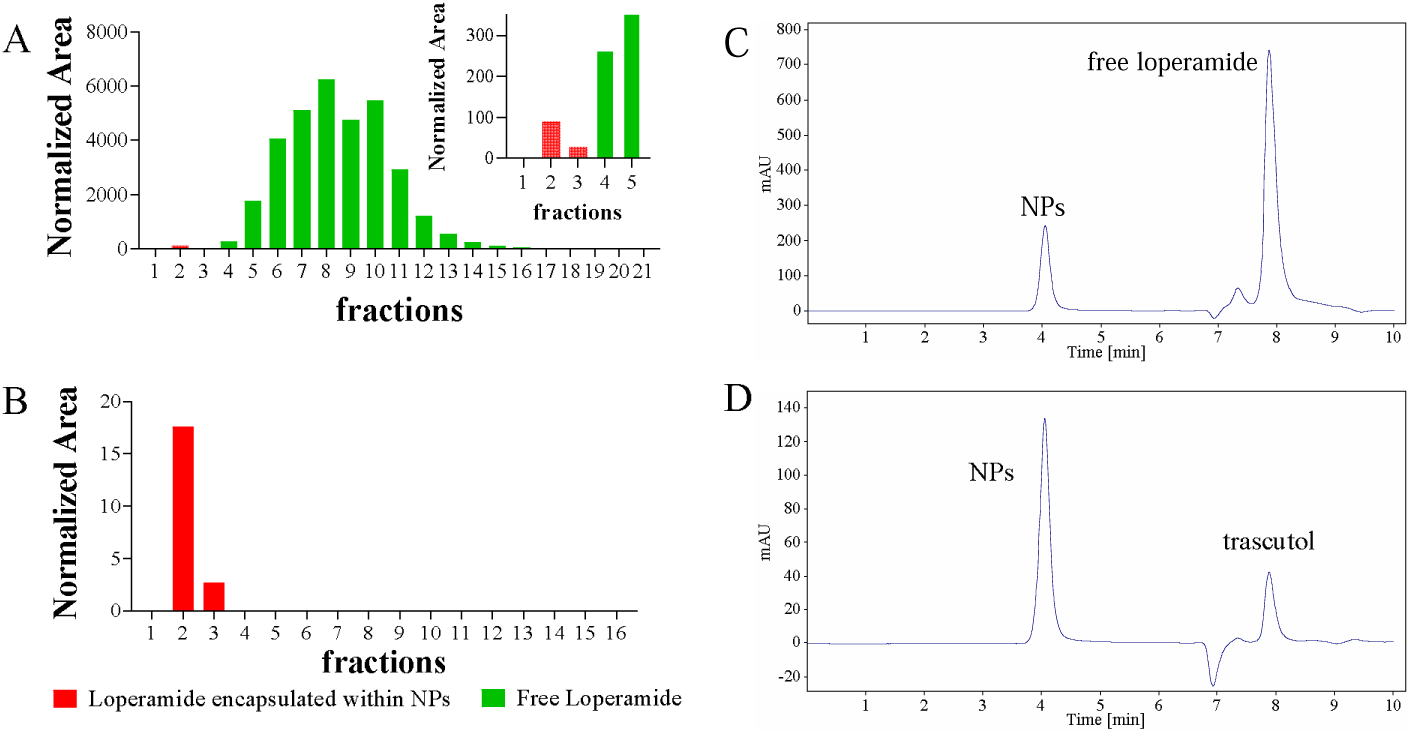
Fraction analysis of samples (A) unwashed SNAP NPs with loperamide and (B) washed (Amicon tubes with molecular weight cut-off, 3 kDa; Millipore) SNAP NPs with loperamide after separation on Zeba Spin Desalting Column. SEC-HPLC analysis for (C) unwashed SNAP NPs with loperamide and (D) washed (Amicon tubes with molecular weight cut-off, 3 kDa; Millipore) SNAP NPs with loperamide.

### 3. Controlled Release, FRET and stability of SNAP NPs

Controlled release of loperamide was studied for PEG-PLA-COOH NPs in PBS at 37°C. It was found that these NPs showed rapid loperamide release, with greater than 50% of drug released within the first 2 hours (**Fig. 4A**).

**Figure 4.**
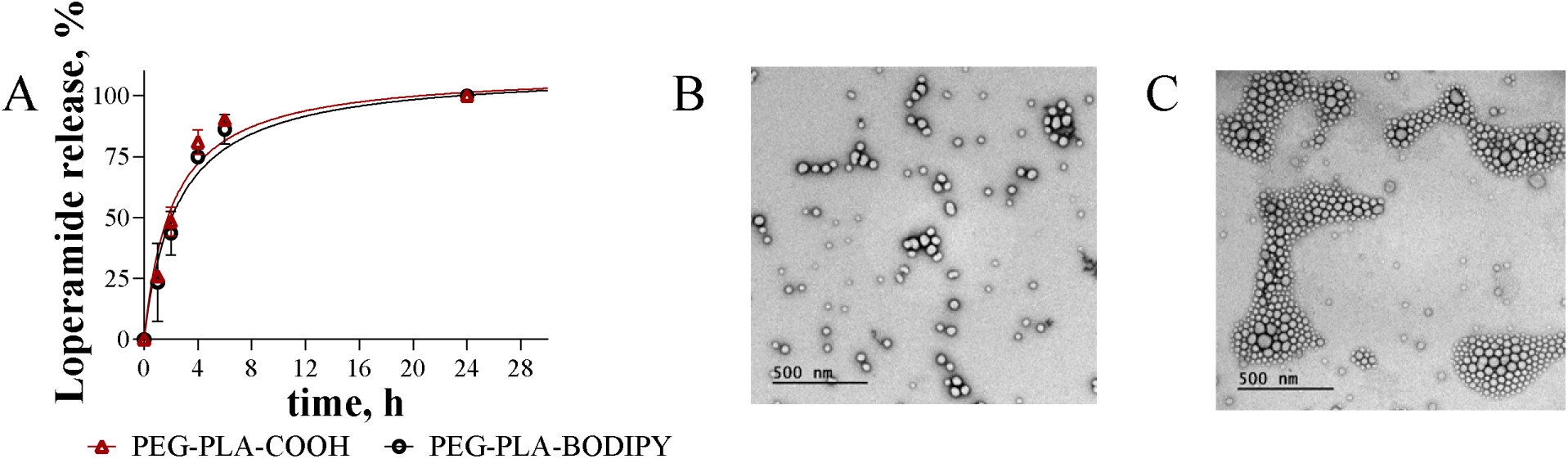
(A) Loperamide release profile of SNAP NPs for different polymers; (B) TEM image of SNAP NPs (PEG-PLA-COOH) with loperamide, (C) TEM image of of SNAP NPs (PEG-PLA-BODIPY) with loperamide.

Drug release may occur via different mechanisms, including rapid diffusion of drug from the hydrophobic PLA core (**Fig. 5A (I)**), diffusion from the PEG layer (**Fig. 5A(II)**), or bulk degradation and disassembly of the NP (**Fig. 5A(III)**). We sought to identify mechanisms of payload incorporation into SNAP NPs using a Fluorescence Resonance Energy Transfer (FRET) technique (**Fig. 5B**). FRET is a process whereby an excited donor molecule (D) transfers energy to a nearby acceptor molecule (A) through a non-radiative interaction (**Fig. 5C**). The strong dependence of FRET on the D−A distance that should be smaller than 10 nanometers (**Fig. 5D**) can provide useful information on drug release and its location within NPs. To study NP stability with FRET, we covalently modified PEG-PLA-COOH with BODIPY FL dye (donor) by EDC/NHS activation and used Rhodamine B (acceptor) as a model payload (**Fig. 5B**). The results showed that these NPs have a clear FRET response (**Fig. 5E and F**). In the presence of the acceptor (Rhodamine B) and donor (BODIPY FL dye) within the intact nanoparticle, the energy was transferred from the excited BODIPY FL dye to Rhodamine B, resulting in donor quenching and acceptor sensitization (**Fig. 5F**). The FRET efficiency was calculated as ∼ 82%. The distance between donor (BODIPY FL) and acceptor (Rhodamine B) chromophores was estimated as 6.2 nm, indicating very close proximity between BODIPY FL dye and Rhodamine B. Therefore, Rhodamine B was encapsulated inside the hydrophobic core of NPs in PLA layer.

**Figure 5.**
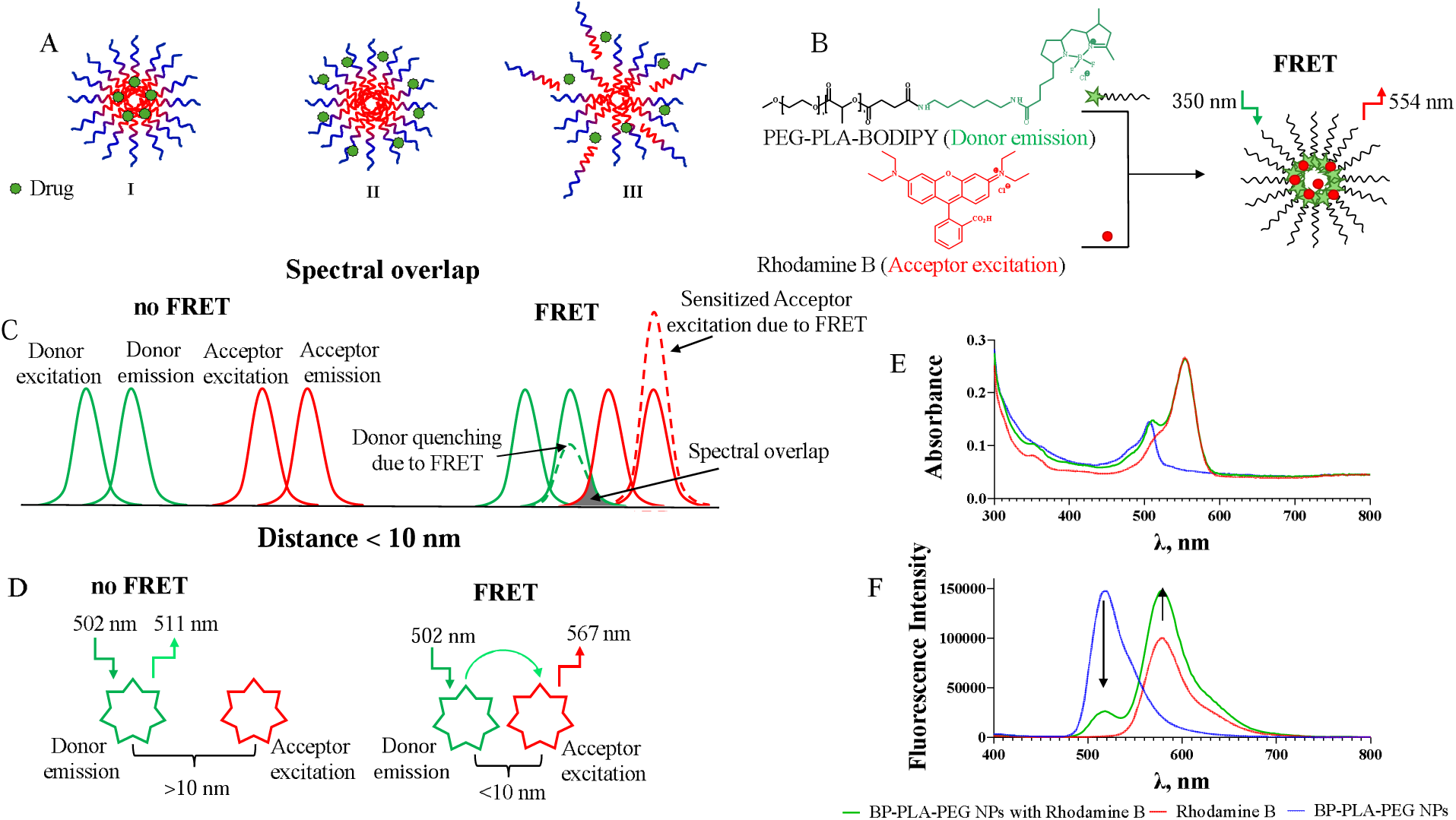
(A) Three possible ways of drug release (I) from hydrophobic PLA core, (II) from PEG shell, (III) NPs degradation. (B) Schematic representation of FRET NPs formulation with Rhodamine B. Explanation of the FRET effect. FRET signal occurs if (C) there is spectral overlap of the donor emission and the acceptor excitation and (d) the distance between the dye molecules is smaller than 10 mn; (E) absorbance of NPs with Rhodamine B, drug-empty NPs and Rhodamine B; (F) emission spectra of NPs based on PEG-PLA-BODIPY with Rhodamine B, drug-empty NPs and Rhodamine B.

To assess the colloidal stability (**Fig. 6A**) of PEG-PLA-C(O)NH-BODIPY NPs loaded with Rhodamine B, we used SEC HPLC and fluorescence spectroscopy. NPs were incubated in 10% FBS in PBS at 37°C for 8 days. Rhodamine B loaded PEG-PLA-C(O)NH-BODIPY NPs were found to be stable for up to 24 hours, as evidenced by a lack of obvious changes in the HPLC chromatograms over this time (**Fig. 6B**). Compared to dye loaded NPs, empty NPs were stable for a shorter period (up to 6 hours) (**Fig. 6C**). A steeper slope on a plot of NPs concentration against time for dye empty NPs also indicates their faster degradation (**Fig. 6C**). Colloidal stability of NPs loaded with Rhodamine B was also demonstrated by fluorescence spectroscopy (**Fig. 6D, E, F**). Figure **6D and E** show that fluorescent intensities at 508 nm that correspond to the NPs increases for 6 hours due to release of Rhodamine B and weakening FRET effect and then decreases after 24 hours due to the degradation of NPs. Figure **6D and F** also show that the light emission at 554 nm (Rhodamine B) decreases over time due to the dye release. Rhodamine B loaded PEG-PLA-C(O)NH-BODIPY NPs was found to have the fastest release kinetics of hydrophilic dye (50% within the first hour) (**Fig. 6G**) compared to lipophilic loperamide (**Fig. 4A**). Therefore, we observe that the model payloads loperamide and rhodamine are likely loaded within the core of the NP, and that NPs loaded with drug are stable over 24 hours. NPs release 98% of drug during this period, after this time in vitro NPs degrade by break down into its constituent materials (**Fig. 6A**).

**Figure 6.**
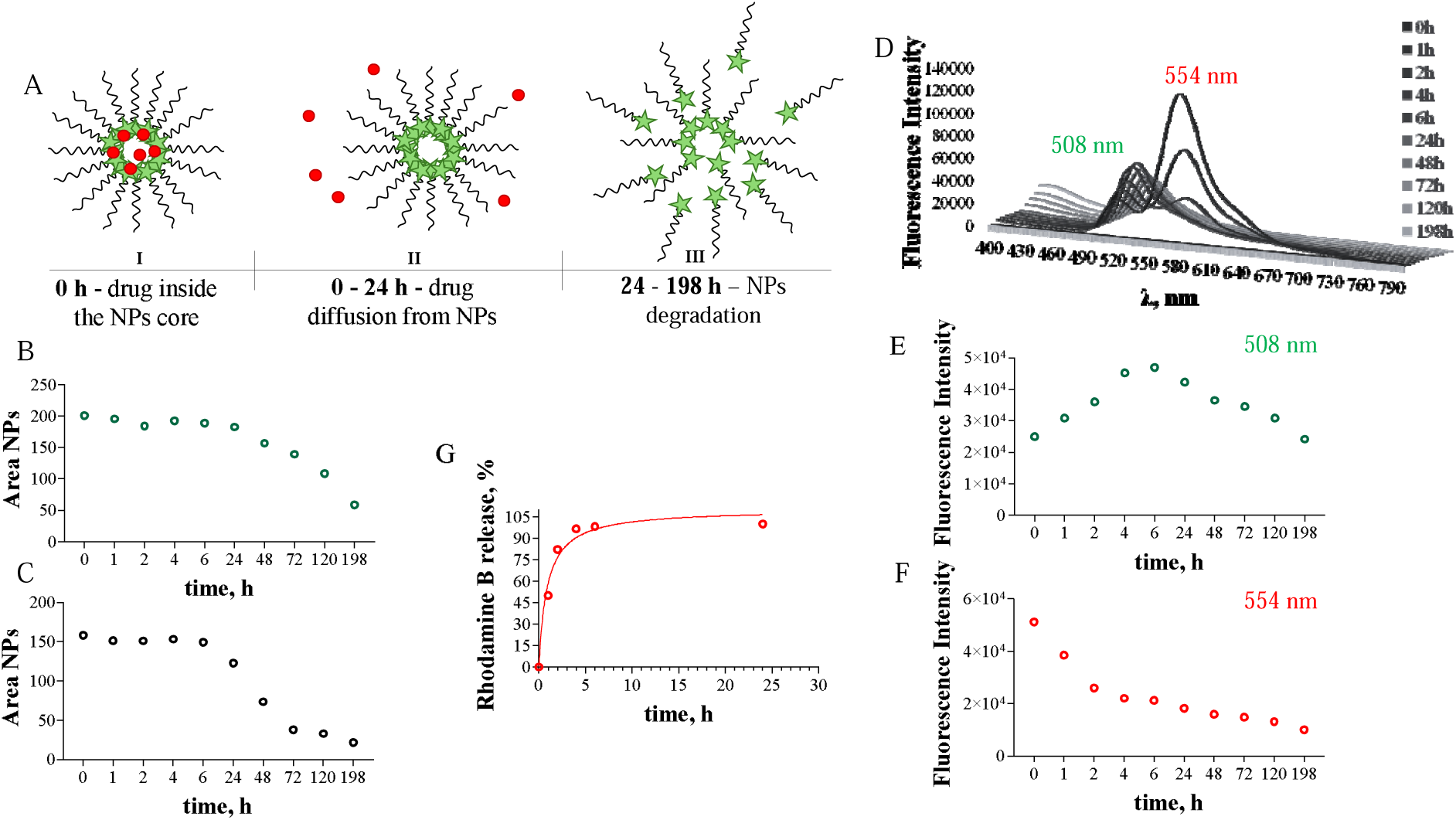
(A) Schematic representation of FRET NPs stability. (B) SEC-HPLC analysis for NPs loaded with Rhodamine B stability. (C) SEC-HPLC analysis for drug-empty NPs stability. (D) The fluorescence intensity changes over time for FRET NPs. (E) A fluorescence curve at 508 nm (NPs). (F) A fluorescence curve at 554 nm (Rhodamine B). (G) Rhodamine B release profile of FRET NPs.

### 4. Impact of BODIPY on NPs properties

To investigate whether the modification of PEG-PLA-COOH with BODIPY FL will impact the NP properties or loperamide loading and release profile, we prepared and characterized PEG-PLA-C(O)NH-BODIPY NPs loaded with this drug. Interestingly, those two formulations (PEG-PLA-COOH with loperamide and PEG-PLA-C(O)NH-BODIPY with loperamide) showed no obvious changes in the particle size, shape, surface charge, drug loading and release kinetics (**Fig. 4 and 7**). Also, no notable changes in particle size, surface charge and release profile were shown for NPs based on PEG-PLA-C(O)NH-BODIPY with loperamide and Rhodamine B except loading (**Fig. 4, 6 and 7**). When NPs based on PEG-PLA-C(O)NH-BODIPY were used for loading two substances with different hydrophobicity (loperamide is more hydrophobic than Rhodamine B), Rhodamine B resulted in lower loading (1.2±0.1 w/w%) compared to loperamide (2.1±0.2 w/w%).

**Figure 7.**
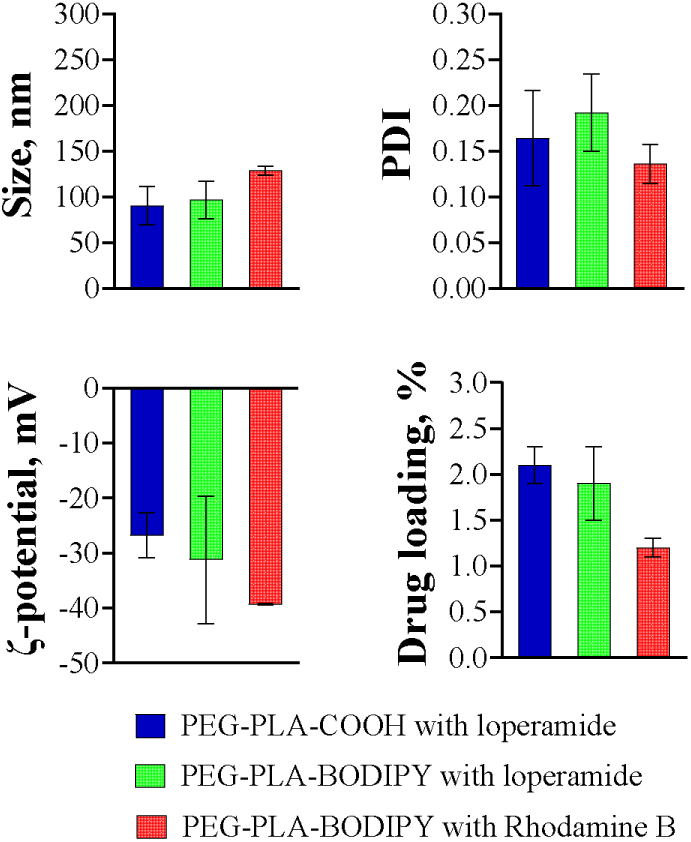
Plot of mean size, PDI, ζ-potential and loperamide and Rhodamine B loading for NPs formed by PEG-PLA-COOH and PEG-PLA-BODIPY FL.

### 5. Effect of drug structure on SNAP NPs properties

To study how the structure of drug influence on NP properties (drug loading, particle size, PDI and ζ-potential), we generated a library of SNAP formulated NPs loaded with drugs of various size and possessing distinct functional groups. As was shown earlier (**Fig. 2B**), the presence of carboxylic groups within polymer (HOOC-PLA-PEG) can significantly influence drug encapsulation by increasing the drug loading from 1.2±0.4% to 2.1±0.2% due to the ability of carboxylic acid fragments to interact with drugs’ functional groups. However, no significant changes in loading were observed for CT179, bortezomib, panobinostat. (**Fig. 8**). CT179 and bortezomib with pyridine and pyrazine fragments can form very strong Lewis acid-base adduct in comparison to Panobinostat, although they yielded similar loading (∼1 w/w%). The lowest loading (less than 0.4 w/w%) was observed for docetaxel, methotrexate and camptothecin, presumably due to their decreased solubility in diethylene glycol monoethyl ether. Diclofenac achieved the smallest size, with its compact molecular structure (shape) likely enabling the highest drug loading (5.1±0.8 w/w%) out of the drugs that we tested (**Fig. 8**). No notable changes in particle size, surface charge and PDI were shown for NPs with different drugs (**SI**).

**Figure 8.**
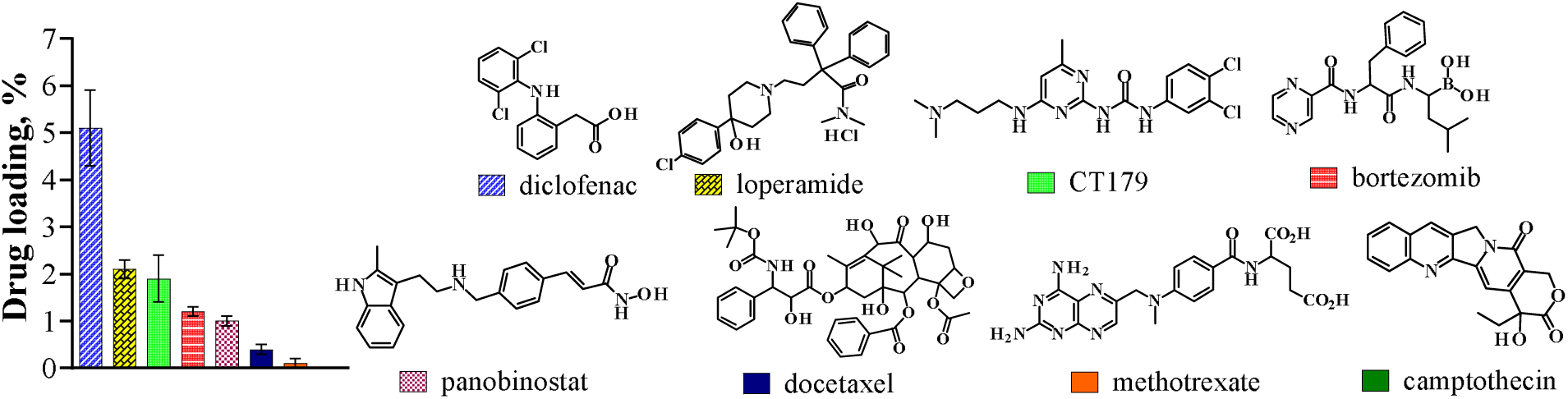
Plot of drugs loading for NPs formed by PEG-PLA-COOH.

### 6. Intravital imaging of fluorescently labeled NPs

To study the delivery of fluorescently labeled NPs to meningeal vessels that supply the brain with blood, we utilized thinned and open skull cranial windows applied to the dorsal aspect of the skull (**Fig. 9A**). NP delivery could be observed by both approaches, with cranial window (**Fig 9C**) yielding a brighter signal than thinned skull (**Fig. 9D**), although NPs were detectable by both methods. To facilitate a direct comparison between the two approaches, the NP signals were normalized. One hour after infusion of PEG-PLA-C(O)NH-BODIPY NPs into cerebrospinal fluid (CSF), via the cisterna magna, we observed NPs within perivascular spaces (PVS) (**Fig. 9C and D**). Notably, NPs appeared as bright and discrete spots or clusters of fluorescence that followed tracks adjacent to blood vessels. Based on the location of the fluorescence signal as well as a characteristic pattern of striation, we are able to conclude that NPs are traveling within the PVS. The vast majority of nanocarriers were seen within 1-7 μm distance from small (sbv), medium (mbv) and large brain vessels (lbv) surface (92%) for more than 3 hours. Occasionally, NPs were noticed inside bifurcations regions of pial vessels and small brain vessels (8%) within 20 μm to the wall surface (**Fig. 9B**).

**Figure 9.**
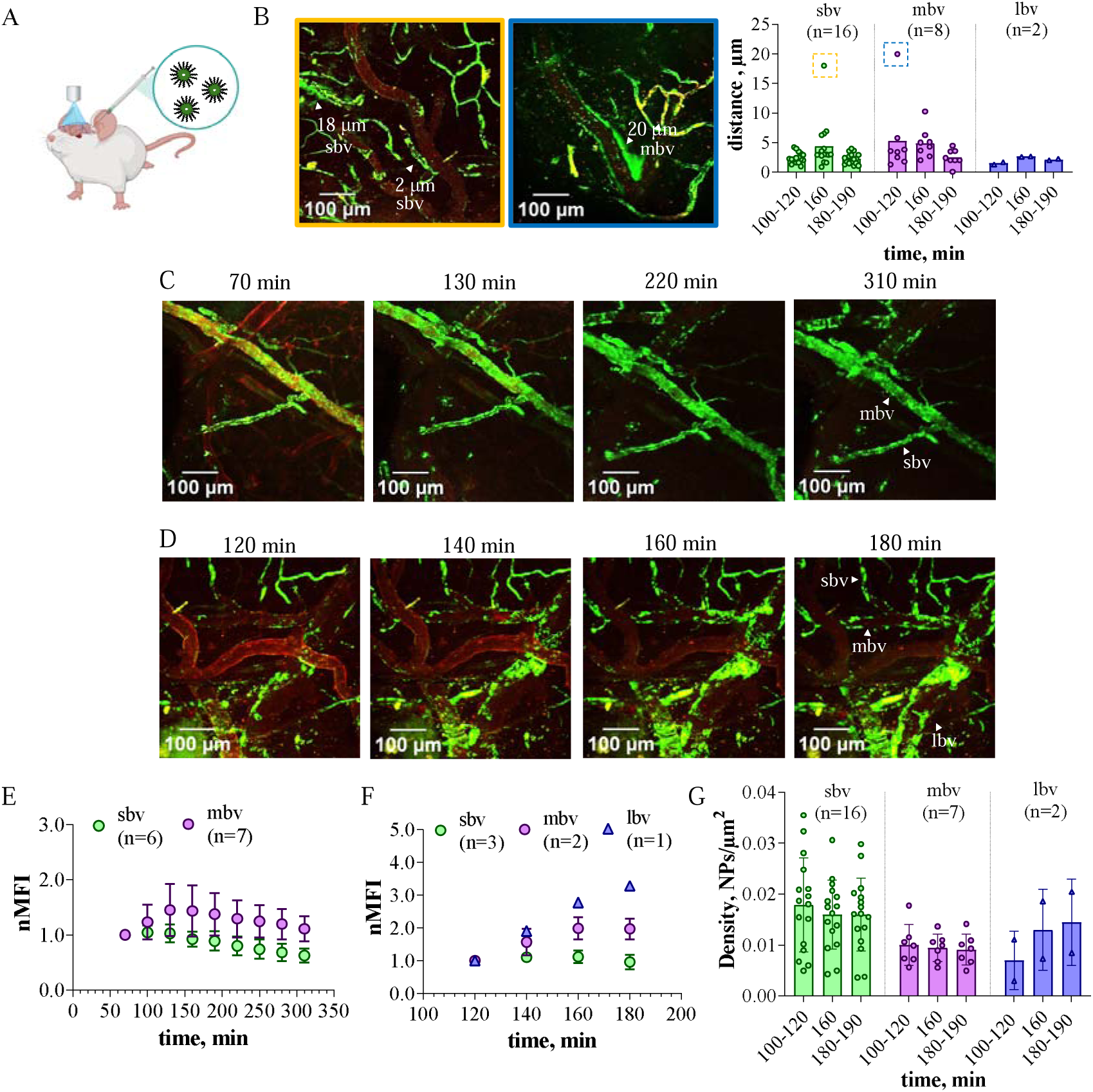
(A) Intravital imaging of NPs (green) based on PEG-PLA-C(O)NH-BODIPY in live mice through a thinned skull (C) and open skull (D) cranial window after cisterna magna injection. (B) The distance from the vessel wall surface occupied by NPs - post-injection time curve of NPs. (B, C, D) Mouse brain vasculature network was labeled by CD31 (red) through tail vein injection. (E) The in vivo normalized median fluorescence intensity (nMFI) - post-injection time curve of NPs for small brain vessels (sbv (<13 μm)) and medium brain vessels (mbv (13-50 μm)). (F) The in vivo normalized median fluorescence intensity (nMFI) - post-injection time curve of NPs for small brain vessels (sbv (<13 μm)), medium brain vessels (mbv (13-50 μm)) and large brain vessels (lbv (>50 μm)). (G) The density of NPs - post-injection time curve of NPs for small brain vessels (sbv (<13 μm)), medium brain vessels (mbv (13-50 μm)) and large brain vessels (lbv (>50 μm)).

To examine the kinetics of NPs distribution we next captured a series of images at regular intervals over 5 hours, focusing especially on medium (13-50 μm) and small (<13 μm) vessels (**Fig. 9C**), and over 3 hours for large (>50 μm) blood vessels of the brain (**Fig. 9D**). Intriguingly, we identified a difference in peak accumulation (C_max_) vs clearance of NPs as a function of vessel size. We therefore performed an additional analysis in which NP delivery kinetics were tracked as a function of vessel diameter. For small (<13 μm) vessels time to maximum concentration (T_max_) was 100-140 min, for medium (13-50 μm) – 140-160 min and for large (>50 μm) was more than 180 min (**Fig. 9E and F**). At 100-120 min post-injection, the highest density of NPs was at walls of small brain vessels, with decreasing along medium to large pial vessels (**Fig. 9G**). However, after 180-190 min of NPs injection, the density of NPs within PVS of large brain vessels increased substantially and became equal to density of NPs at walls of small pial vessels, meanwhile the NPs density at surface of small and medium brain vessels decreased insignificantly overtime (**Fig. 9G**). Therefore, the kinetics of NPs accumulation were fastest for small and slowest for large pial vessels.

To study nanocarrier movement dynamics, NPs were manually tracked at 60 min and 160 min post-injection for 1 min of continuous video within PVS of medium and small vessels (**Fig. 10A**) (**Supplementary Movie 1**). To understand the general pattern of nanoparticle movement, the individual traces (paths) were aligned to the point of origin (**Fig. 11B**). Tracking analysis demonstrated that motility of NPs was higher at medium pial vessels compared to the small one within the first 60 min after administration followed by a significant decrease in motility after 160 min (**Fig. 10B**). Despite differences in vessel size, net displacement of NPs (i.e., distance between the start and end points) did not significantly differ between primary and daughter vessels (**Fig. 10C**). However, for both small and medium vessels, it was ∼ 54% shorter after 160 min compared to 60 min post-injection (**Fig. 10C**). This was consistent with the average velocity of NPs being fastest at 60 min post-injection and slowest at 160 min (**Fig. 10D**). We also observed that over time some of the NPs were trapped within PVS on the vessels wall. (**Supplementary Movie 2**). Taken in sum, these results identify vessel size-dependent differences in motility of NPs along the PVS after direct infusion into cerebrospinal fluid.

**Figure 10.**
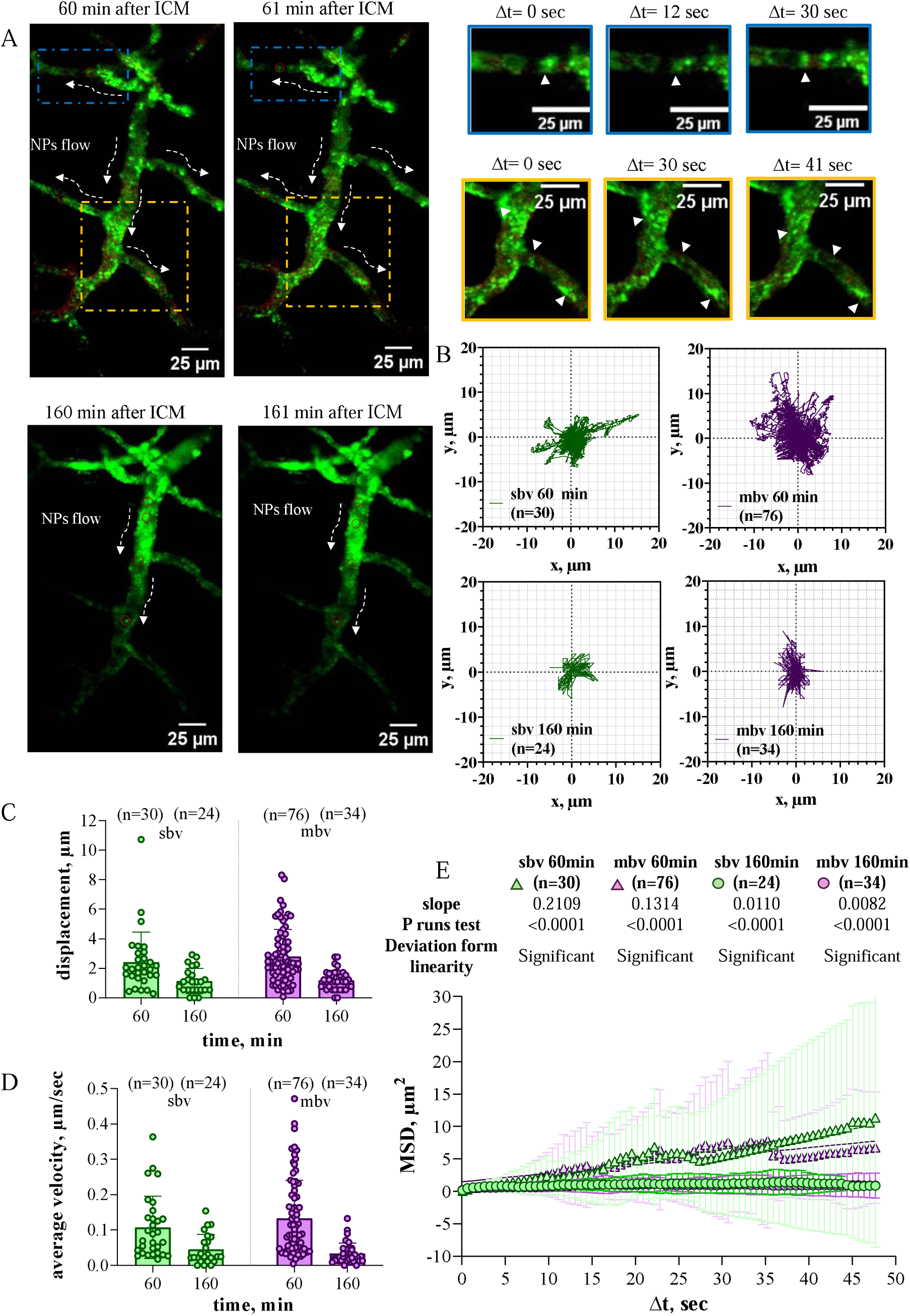
(A) Intravital imaging representing movement of NPs (green) based on PEG-PLA-C(O)NH-BODIPY in live mice through a thinned skull cranial window after cisterna magna injection. Mouse brain vasculature network was labeled by CD31 (red) through tail vein injection. (B) Pattern of NPs movement, aligned to the point of origin 60 min and 160 min post-injection. (C) 60 sec displacement of NPs - post-injection time curve of NPs for small brain vessels (sbv (<13 μm)) and medium brain vessels (mbv (13-50 μm)). (D) Average velocity of NPs - post-injection time curve of NPs for small brain vessels (sbv (<13 μm)) and medium brain vessels (mbv (13-50 μm)). (E) MSD analysis of trajectories of individual NPs.

To assess whether PVS trafficking of NPs is driven by stochastic molecular motion (i.e., diffusion) or bulk flow (i.e., convection), we performed a mean squared displacement (MSD) analysis of trajectories of individual NPs (**Fig. 10E**). It was found that nanoparticle movement deviates significantly from linearity for 60 min compared to 160 min post-injection despite the vessel size (**Fig. 10E**). This indicates that NPs transport within the PVS is not solely governed by simple diffusion but rather a coordinated form of trafficking within PVS, occurring both within the smallest - daughter blood vessels and larger – primary vessels.

## Discussion

The one-step nanoprecipitation technique is a promising approach for creating drug-delivery vehicles based on NPs. Fessi and co-workers were the first to develop and patent solvent displacement method for formulation of nanocarriers [10,11]. This technique reduces time, simplifies the process and improves efficiency [12, 13]. Rapid formulation of nanoparticles allows quickly generate and test various nanocarriers in vivo, accelerating the process of optimizing nanoparticles properties like size, shape, surface chemistry, drug loading and release profile for specific therapeutic applications. Several techniques like solvent evaporation, nanoprecipitation, salting-out, and emulsification can be used to create drug loaded nanocarriers with desired properties to achieve specific therapeutic effects [14, 15]. However, not all nanoparticle formulation techniques allow for the easy and rapid fabrication of nanocarriers with controlled properties. Different approaches have varying levels of control over size, shape, and composition, and some can be more complex, time-consuming, or require specialized equipment. Importantly, compared to nanoprecipitation, emulsion-based systems have faced serious obstacles regarding challenges in removing residual solvents, limitations in encapsulating of certain drug types and manufacturing scale up. Another technique – solvent evaporation method that can be used for NPs formulation also present several disadvantages including the cross-contamination, difficulty in controlling the evaporation rate and inefficient of scale up due to energy requirements and potential instability of heat-sensitive compounds, these problems can be solved using nanoprecipitation or one-pot approaches. However, despite the potential advantages of one-pot approaches described early, controlling NPs size, surface charge and morphology affecting drug loading capacity, release kinetics and in vivo behavior remain a challenge. Optimization existing formulation approaches or development new ones are necessary to create drug loaded nanocarriers with controlled properties.

In this study we described a new one-pot technique to NP formulation, which we term Solvent-free Nanoparticle Assembly Protocol (SNAP). SNAP enables rapid and reproducible core-shell NP formulation based on amphiphilic copolymers (PLA-PEG and PCL-PEG), offering control over drug release and loading, size, shape, and surface properties of nanocarriers. To create NPs via SNAP, we used diethylene glycol monoethyl ether for dissolving amphiphilic copolymers. Diethylene glycol monoethyl ether is a versatile excipient used in pharmaceutical and cosmetic formulations to enhance drug delivery. It helps dissolve poorly soluble drugs, acts as a solvent in the preparation of NPs, for example, producing smaller liposomes compared to other organic solvents [16], and improves drug penetration into the skin or other tissues. Diethylene glycol monoethyl ether that listed in the FDA’s IID is considered safe due to its well-established safety profile, supported by nonclinical data and extensive use as a vehicle and solvent across various administration routes (the specified limits are up to 49.9% for topical use and up to 5% for transdermal use) [17]. Based on the safety profile of diethylene glycol monoethyl ether, we do not need to remove it prior to use in vivo, that can be additional advantages of SNAP due to simplify the overall nanoparticle formulation, making it faster and more cost-effective.

The size and surface charge (ζ-potential) are major parameters of drug-delivery vehicles affecting the carrier’s stability, release profile, cellular uptake, tissue penetration, biodistribution, circulation time and clearance [18, 19]. Importantly, for NPs composed of a variety of biomaterials, including polyesters or biologically derived constituents, the size, surface properties and drug loading of NPs may be controlled by changing of molecular weight or concentration [20–24]. To evaluate the drug-loading capacity of SNAP-formulated NPs, we used loperamide as a model compound for CNS drug delivery. When loperamide is administered in free form, it is unable to bypass the BBB, however, when it is administered directly to the CNS or has been incorporated into NPs that have BBB crossing capability, antinociceptive effects are observed. In this study, we formulated NPs loaded with loperamide via SNAP utilizing amphiphilic polymers, either polylactic acid (PLA) or polycaprolactone (PCL). Loperamide loaded NPs formulated with PCL have the size in the range of 43.8±1.5 to 190.8±6.5 nm and lower value of ζ-potential −26.3±0.7mV. In contrast, PLA can form NPs in the 76.9±2.6 to 144.6±8.0 nm range with lowest ζ-potential −38.6±3.4 mV. Therefore, we can conclude that PLA NPs can form more stable (ζ-potential < −35 mV, pH=7) loperamide loaded NPs with the narrow size distribution (PDI < 0.2) and small size. The small size of NPs generated by SNAP may enhance their ability to penetrate tissues and cross biological barriers. Despite the advantages of these nanocarriers, low drug loading capacity could remain an issue, significantly reducing their effectiveness. In this study we showed that loperamide loading for SNAP NPs did not exceed 1 w/w% (encapsulation efficiency 0.5 w/w %). Hydrophobic interactions for SNAP NPs between a drug (loperamide) and a PLA (poly(lactic acid)) polymer could be insufficient to retain the drug within the PLA core. To increase the drug loading different functional groups on polymers, like amino, carboxyl, or thiol groups, can be used to attract and bind drugs, improving drug encapsulation [25]. We applied this approach to our platform. It was shown that the presence of carboxylic groups (COOH) within the PLA structure facilitates the incorporation of drugs with basic groups (loperamide, CT179, bortezomib, panobinostat) into the core of the NPs, increasing encapsulation efficiency due to Lewis acid (PLA-COOH) - base (drug) interaction. The loading for these drugs varies from 0.1±0.1 to 2.1±0.2% (encapsulation efficiency 0.6±0.1-11.1±1.4%), which is typical for small, micellar-like NPs based on PLA-PEG [24, 26]. However, not only the presence of different functional groups on polymers influences the drug loading, the size and shape of the drug could significantly affect the drug encapsulation as well. Small drugs are more likely to fit within limited internal space (core) of the small carriers (with size less than 100 nm). We demonstrate that SNAP NPs can encapsulate small drugs like diclofenac with DL 5.1±0.8% (encapsulation efficiency 12.9±5.8%), making this platform more suitable for delivering small hydrophobic drug with compact shape. Despite that the loading of bortezomib [27], and diclofenac [28] into other polymeric NPs can be more than 9%, the size of some of the nanocarriers can exceed size of NPs generated by SNAP.

Drug release is an important characteristic that impacts the safety and overall therapeutic effectiveness of the drug. This parameter may be controlled by engineering the core and shell of NPs. The core can be used to encapsulate the drug and keep it inside the NPs due to van der Waals forces, hydrophobic interaction and/or hydrogen bonding, while the shell acts as a barrier, controlling the rate at which the drug is released. There are two major pathways of controlled drug release from nanocarriers: (I) drug diffusion from intact NPs through both core and shell layers and (II) release due to bulk degradation. To estimate the location of drug (core or shell) within NPs based on PEG-PLA-C(O)NH-BODIPY we used a FRET technique. FRET relies on the transfer of energy between two fluorophores, a donor (PEG-PLA-C(O)NH-BODIPY) and an acceptor (Rhodamine B - model drug). After the donor excitation we observed its energy transfer to the acceptor indicating close proximity (6.2 nm) between BODIPY and Rhodamine B. Location of the drug within NPs core and dense PEG shell can enhance controlled release. We demonstrate that SNAP NPs based on PLA-PEG release drug from the hydrophobic core of intact NPs through shell layer within the 24 h. After this time, the NP experiences bulk degradation and eventual disassembly. In this study, we showed that the degradation kinetics of SNAP NPs depends on the presence of drugs inside the core. Drug molecules likely slow down the degradation rate of nanocarriers due to slower penetration of water into the polymer matrix or due to the interaction (hydrophobic, van der Waals, hydrogen bonding) with polymer chains forming the core of NPs.

To estimate SNAP NPs behavior in vivo we used intravital microscopy (IVIM) that enables real-time observation of cells, tissues, nanocarriers within a living animal. Understanding the relationship between nanoparticles properties and their behavior within the complex biological system is crucial for successful translation of nanoparticle-based therapies. Fluorescently labeled NPs based on PLA-PEG were injected intravenously, and the liver was imaged through an abdominal window. After intravenous injection, colloids in the bloodstream will distribute throughout the body, with NPs larger than 60-80 nm more likely to be deposited in the liver (from 30% to 99% of total administered dose) [29, 30], whereas NPs smaller than 20 nm experience primarily renal clearance [31]. Surface chemistry is another consideration. Positively charged NPs are more likely taken up by hepatocytes, while negatively charged NPs are preferentially captured by Kupffer cells and liver sinusoidal endothelial cells. Park at al. showed the highest concentration (98%) of negatively charged biodegradable poly(lactic-co-glycolic acid) (PLGA) particles were taken up by Kupffer cells [32]. Excessive NP accumulation in the liver can reduce their delivery to targeted tissues, hindering the effectiveness of certain therapeutic and imaging applications. One of the possible approaches for circumventing this natural barriers like liver is modification of nanoparticle surface charge to neutral or slightly negative by coating the surface with polymers like PEG (polyethylene glycol). PEGylation makes particles less recognizable by liver cells and immune system and allows them to circulate longer. Gref at al. showed that 66% of noncoated PLGA NPs were removed by the liver 5 min after injection while only 30% of PEG-coated NPs were captured by the liver 2 hours after administration and still present even after 5 hours. [33]. This data agrees with our results (**Fig. S11**), showing by IVIM that SNAP NPs based on PLA-PEG are still detectable in the liver 4 hours after intravenous injection. The accumulation of NPs occurs during the first hour of introduction, then clearance mechanisms presumably mononuclear phagocyte system (Kupffer cells) slowly begins to remove them from the liver. Extending the circulation lifetime (more than 4 hours) of SNAP NPs in the bloodstream of large (>50 μm) vessels were also observed through the cranial window after intravenous administration. Thus, we demonstrated that PLA-PEG NPs have longer circulation time in blood flow that allows the drug delivery vehicle based on SNAP NPs to reach the target tissue more effectively before being cleared by the body.

Intrathecal injection (IT) is one of the methods that can be used to circumvent the blood-brain barrier and deliver NPs directly to the central nervous system. After injection into cerebrospinal fluid, nanocarriers can enter perivascular space of pial vessels separated from the subarachnoid space by a thin meningeal sheet [34]. Bedussi at al. showed that 1 μm fluorescent labeled microspheres after cisterna magna administration moved preferentially in the perivascular space of arteries within a 20-40 μm range next to the vessels rather than within the adjacent subarachnoid space or the space surrounding veins [35, 36]. This corresponds well with our data. In our study, we demonstrated extensive presents of PLA-PEG NPs close to the small and medium vessels and sometimes around the large vessels within 1-20 μm range to the vessel surface. A previous study showed that movement of microspheres, generated by the heartbeat, along perivascular space of arteries was pulsatile and in the same direction as blood flow [35, 36]. In the present study, we confirmed the flow of PLA-PEG NPs from large vessels to smaller ones. We also showed that the movement of NPs is not solely driven by simple diffusion, they are actively transported within the PVS. Based on flow direction of PLA-PEG NPs within PVS we could assume that SNAP NPs nanocarriers occupied perivascular space of arteries. We demonstrate that NP transport within PVS is influenced by the size of the blood vessels. For the small vessels (< 13 μm) NPs reach their maximum concentration (C_max_) in the PVS relatively quickly within 100-140 min (T_max_). For medium vessels (13-50 μm) T_max_ is slightly longer (140-160 min). For large vessels (> 50 μm) T_max_ is the longest, taking over 180 min to reach C_max_. Smets at al. showed that the PVS around arteries exceed the perivenous spaces and creates a low resistance pathway for CSF flow [34]. The size of PVS increased by increasing the diameter of the vessels [34, 37]. In the present study, we also confirmed that NPs are able to move more effectively within PVS of larger pial vessels at least during the first hour after administration. We also observed that PLA-PEG NPs can be trapped overtime on the vessel surface (particularly near the bifurcation areas) as a result decreasing the displacement (distance from the initial (starting) point to the final (ending) point) and slowing down average velocity of nanocarriers within PVS. Based on this data we speculate that the small size of PVS around capillaries and their branching geometry facilitate NPs to interact with vessel surface and be deposited on it more rapidly (T_max_ - 100-140 min). While bigger pial vessels due to simpler geometry, larger PVS and low resistance pathway for CSF flow within PVS may require more time for NPs to be trapped on the vessel wall (T_max_ >180 min). Deposition of NPs on the wall surface of big pial vessels could slow down their movement overtime leading to transfer less NPs to smaller daughter vessels. It was confirmed by showing that the density of NPs within PVS of large brain vessels increased significantly and eventually reached a density comparable to that observed at the walls of smaller pial vessels. Thus, in this study we have demonstrated that biodegradable PLA-PEG NPs with the size around 100 nm are able to move along PVS of brain vessels and stay in the SAS more than 5 hours. Understanding how NPs behave within the living brain is essential for developing drug delivery systems for treatment of neurological disorders. This knowledge can be useful in designing more effective nanocarriers that can reach specific brain regions and deliver therapeutic agents directly to the target cells for maximizing therapeutic efficacy and minimizing toxic side effects.

## CONCLUSION

In this study, a novel Solvent-free Nanoparticle Assembly Protocol (SNAP) was developed for facile NP formulation. SNAP is a very simple and quick method for making NPs with well-controlled particle size, size distribution, surface properties and drug loading. We demonstrated that nanocarriers based on PLA-PEG with the size around 100 nm can potentially be used as a platform for encapsulation and delivery small hydrophobic drugs. Colloidal stability for 24 hours and controlled drug release of SNAP NPs could prevent high initial drug concentrations in vivo and provide a more sustained therapeutic effect, given excellent colloidal stability. In vivo studies have demonstrated that SNAP NPs have longer circulation time in blood flow, slow accumulation and clearance within perivascular space of big arterials. Longer circulation time in blood and retention of NPs within SAS may increase their delivery efficiency and offer more effective and prolonged therapeutic effects of encapsulated drugs. In sum, these studies describe a new approach for NP formulation that can be used for purposes ranging from drug delivery to imaging.

## METHODS

### Materials

PLA-PEG, PEG-PLA-COOH and PCL-PEG (PolySciTech), Invitrogen UltraPure DNase/RNase-Free distilled water (Fisher Scientific), diethylene glycol monoethyl ether (>99.0%, TCI America), Sodium phosphate, monobasic dihydrate (HPLC garde, Thermo Scientific), loperamide (>98.0%, TCI America), diclofenac (98%, Thermo Scientific), CT179 (Curtana Pharmaceuticals), bortezomib (PS-341, APExBIO), Panobinostat (LBH589, APExBIO), docetaxel (99%, Thermo Scientific), methotrexate (Thermo Scientific), camtothecin (≥98% (HPLC), Enzo), rhodamine B (MilliporeSigma), EDC (Premium grade, Thermo Scientific), sulfo-NHS (Premium grade, Thermo Scientific), triethylamine (Thermo Scientific), dimethyl sulfoxide (≥98%, Sigma Aldrich), Sodium chloride (≥99%, Sigma Aldrich), Cyanine 5 amine (Lumiprobe), BDP FL amine (97%, BroadPharm), Fetal Bovine Serum (FBS) (Cytiva), gibco DPBS (1x, Thermo Scientific), Alexa Fluor 647 anti-mouse CD31 (BioLegent), size-exclusion column Agilent BIO SEC-5, 100A, TSKgel ODS-0120T C-18 column, 15cm×4.6mm, 5µm, Zeba Spin Desalting Columns, 7K MWCO (Thermo Scientific), Slide-A-Lyzer G3 Dialysis Cassettes, 20K and 3.5K MWCO, Amicon Ultra Centrifugal Filter, 3 kDa MWCO 15mL (MilliporeSigma), microplate reader (Tecan, Spark Multimode Microplate Reader), dynamic light scattering (Malvern Panalytical, Zetasizer Pro), transmission electron microscopy (FEI Tecnai Spirit 12), Allegra X-30 Series Benchtop Centrifuges (Beckman Coulter), 1220 Infinity LC System HPLC, Bruker AVANCE III HD 500 MHz NMR, Scintica IVIM confocal microscope.

### Preparation of drug empty SNAP NPs

To prepare drug empty PLA-PEG and PCL-PEG NPs we employed nanoprecipitation techniques. PLA-PEG and PCL-PEG were dissolved separately in diethylene glycol monoethyl ether at concentrations 25 mg/mL, 50mg/mL and 100mg/mL. Then solutions of polymers (PLA-PEG and PCL-PEG) were sonicated in sonication bath (1 min), after this placed in water bath and heated to 70°C. The polymer solutions were then quickly nanoprecipitated in 5 mL of cold water. Upon nanoprecipitation, NPs formed instantly and were kept for 5 min at 1000 r.p.m. stirring at room temperature. The NPs were then washed three times with cold water using Amicon® Ultra Centrifugal Filter (molecular weight cut-off, 3 kDa). The NPs were used fresh or kept at − 20°C to use later for various in vitro and in vivo studies. Every NPs formulation was repeated 3 times for each condition.

### Preparation of drug loaded SNAP NPs

To prepare drug loaded NPs PLA-PEG, PCL-PEG and HOOC-PLA-PEG were dissolved separately in diethylene glycol monoethyl ether at concentrations 25 mg/mL, 50mg/mL and 75mg/mL. The polymers were mixed with 1 mg of drugs (loperamide, diclofenac, CT179, bortezomib, panobinostat, docetaxel, methotrexate and camtothecin) in a small glass vial. Then the solutions of polymers with drugs were quickly nanoprecipitated in 5 mL of cold water and were kept for 5 min at 1000 r.p.m. stirring at room temperature. The NPs were then washed three times with cold water using Amicon® Ultra Centrifugal Filter (molecular weight cut-off - 3 kDa) to remove free drugs. The NPs were used fresh or kept at − 20°C to use later for various in vitro and in vivo studies. The loperamide-loaded NPs were subjected to Zeba Desalting column and SEC HPLC to check for unencapsulated drug molecules. Every NPs formulation was repeated 3 times for each condition.

### Zeba Spin Desalting Column

To assess the unencapsulated drugs, washed and unwashed loperamide-loaded NPs were run through Zeba Spin Desalting Columns (molecular weight cut-off - 7 kDa). Desalting columns were equilibrated with 150mM NaCl at pH 7.0. After Zeba Spin column preparation, 2 mL of washed and unwashed loperamide-loaded NPs were slowly applied to the center of the compacted resin bed. To collect samples column was centrifuged at 1500 × g for 2 minutes. This step was repeated 21 times by using 2 mL of equilibration buffer. All collected 21 fractions were analyzed using UV-vis, DLS and C18 HPLC (Agilent 1220 Infinity LC System HPLC, TSKgel ODS-0120T C-18 column, 15cm×4.6mm, 5µm).

### High Performance Liquid Size-Exclusion Chromatography (HPLC SEC)

For chromatographic separation of loperamide loaded NPs and unencapsulated loperamide a size-exclusion column Agilent BIO SEC-5, 100A was applied, using 1220 Infinity LC System HPLC with a diode array UV/VIS detector. The applied mobile phase was 150 mM NaH_2_PO_4_, the injection volume 20ul, the flow rate 0.3 mL/min, and the duration of chromatographic run 10 minutes, λ=220nm. ChemStation with Agilent OpenLAB software was used for controlling the system operation and data analysis.

### Drug loading

To calculate the drug loading (DL, %) several concentrations of drugs (loperamide, diclofenac, CT179, bortezomib, panobinostat, docetaxel, methotrexate and camtothecin) (100 µg/mL to 5 µg/mL) for standard curve were prepared in buffer solutions acetonitrile/40 mM NaH_2_PO_4_ (diclofenac, loperamide, bortezomib, camptothecin, docetaxel, CT179 - 40:60, v/v, panobinostat - 20:80, v/v, methotrexate – 10:90, v/v). 200 µL aliquots of nanoparticles were put in a pre-weighed tube and lyophilized. After lyophilization nanoparticles were dissolved in 200 µL of buffer solutions. The solution of NPs was analyzed by reverse-phase HPLC method (Agilent 1220 Infinity LC System HPLC, TSKgel ODS-0120T C-18 column, 15cm×4.6mm, 5µm). The mobile phase was acetonitrile/40 mM NaH_2_PO_4_ (diclofenac, loperamide, bortezomib, camptothecin, docetaxel, CT179 - 40:60, v/v, panobinostat - 20:80, v/v, methotrexate – 10:90, v/v), injection volume was 20 µL and the column temperature maintained at 30°C. The analysis was performed at a flow rate of 1.0 mL/min for diclofenac, loperamide, bortezomib, camptothecin, panobinostat, methotrexate, CT179 and 1.7 mL/min for docetaxel –with the UV detector at 220 nm. Drug loading (DL %) was calculated using formula below: Drug loading (DL%) = (W_drug_/W_NPs_) ×100 % where W_drug_ is the mass of drug measured and W_NPs_ is the mass of nanoparticles. Each experiment was repeated three times.

### Encapsulation efficiency

To determine the encapsulation efficiency of loperamide in nanoparticles, 50 µL aliquot of drug loaded NPs was mixed with 50 µL of acetonitrile/40 mM NaH_2_PO_4_ (diclofenac, loperamide, bortezomib, camptothecin, docetaxel, CT179 - 40:60, v/v, panobinostat - 20:80, v/v, methotrexate – 10:90, v/v). The solution was analysed by reverse-phase HPLC method (Agilent 1220 Infinity LC System HPLC, TSKgel ODS-0120T C-18 column, 15cm×4.6mm, 5µm). The mobile phase was acetonitrile/40 mM NaH_2_PO_4_ (diclofenac, loperamide, bortezomib, camptothecin, docetaxel, CT179 - 40:60, v/v, panobinostat - 20:80, v/v, methotrexate – 10:90, v/v), injection volume was 20 µL and the column temperature maintained at 30°C. The analysis was performed at a flow rate of 1.0 mL/min for diclofenac, loperamide, bortezomib, camptothecin, panobinostat, methotrexate, CT179 and 1.7 mL/min for docetaxel –with the UV detector at 220 nm. Encapsulation efficiency (EE %) was calculated using below formula: Encapsulation efficiency (EE %) = (W_t_/W_i_) ×100 % where W_t._ is the total amount of drug in the nanoparticles and W_i_ is the total quantity of drug added initially during preparation. Each experiment was repeated three times.

### Drug release

To perform drug release experiments 1000 µL solutions of nanoparticles were mixed with 1000 µL of PBS at pH = 7.4 Then 2000 µL solutions of loperamide loaded nanoparticles were transferred to a dialysis cassette (Slide-A-Lyzer G3, 2K MWCO, 3mL). Dialysis cassette was immersed in PBS at pH = 7.4 and incubated at 37 °C. At pre-determined time intervals 0h, 1 h, 2 h, 4 h, 6 h, 24 h and 48 h 50 µL sample was taken from dialysis cassette. The amount of loperamide released from the dialysis was analyzed by reverse-phase HPLC method (Agilent 1220 Infinity LC System HPLC, TSKgel ODS-0120T C-18 column, 15cm×4.6mm, 5µm). The mobile phase was acetonitrile/40 mM NaH_2_PO_4_ (40:60, v/v), injection volume was 20 µL and the column temperature maintained at 30°C. The analysis was performed at a flow rate of 1.0 mL/min with the UV detector at 220 nm. Each experiment was repeated three times.

### NPs characterization

To prepare NPs for Dynamic Light Scattering (DLS) and Electrophoretic Light Scattering (ELS) 100 µL of aliquot of NPs was mixed with 2000 µL of DI water. NPs were characterized by assessing their hydrodynamic diameter, PDI and zeta potential using the Malvern Panalytical Zetasizer Pro. To prepare particles for transmission electron microscopy 5 µL of sample was placed onto a formvar/carbon 200 mesh copper grid for 30 seconds, after which 5 µL of uranyl acetate was placed for negative staining. A transmission electron microscope Tecnai Spirit 12 microscope (FEI Company) was used to assess the morphology and shape of the NPs.

### NPs stability

To perform drug release experiments and mimic in vivo conditions 1000 µL solutions of nanoparticles were mixed with 1000 µL of 10% FBS in PBS. Then 2000 µL solutions of NPs were transferred to a dialysis cassette (Slide-A-Lyzer G3, 20K MWCO, 3mL) and incubated in 10% FBS in PBS solution at 37 °C. At each time point (0, 1, 2, 4, 8, 12, 24, 72, 120, and 198 h), 50 µL of NP solution was taken from dialysis cassette. For particle analysis SEC HPLC (SEC column Agilent BIO SEC-5, 100A) was used. The analysis was performed at a flow rate of 0.3 mL/min with the UV detector at 495 nm. Rhodamine B NPs and drug empty NPs labeled with BODIPY were used in this study.

### Preparation of PEG-PLA-Cy5 and PEG-PLA-BODIPY

PEG-PLA-Cy5 and PEG-PLA-BODIPY were synthesized by functionalization of the carboxylic acid (−COOH) group of PEG-PLA-COOH by EDC NHS reaction. In brief, 30 mg of PEG-PLA-COOH (MW – 5000:16000 Da) was resuspended into 1000 µL DMSO and put on a magnetic stirrer. Thereafter, 2.9 mg of EDC and 3.3 mg of NHS were dissolved in 300ul 1xPBS and added to PEG-PLA-COOH to react at room temperature for 15 min. 1.6 mg of BODIPY amine and 2.5 mg of Cy5 amine separately were dissolved in 500 µL DMSO and mixed with 4 µL of triethylamine, then dropwise added to activated PEG-PLA-COOH and kept overnight for stirring. PEG-PLA-Cy5 and PEG-PLA-BODIPY were washed multiple times using Amicon® Ultra Centrifugal Filter (molecular weight cut-off, 3 kDa) to remove excess of BODIPY amine and Cy5 amine. Then PEG-PLA-Cy5 and PEG-PLA-BODIPY were lyophilized and kept at − 20°C to use later for in vitro and in vivo studies. PEG-PLA-Cy5 and PEG-PLA-BODIPY were characterized by NMR and UV-vis.

### NPs formulation for FRET system

Rhodamine B loaded PEG-PLA-BODIPY nanoparticles were prepared via a nanoprecipitation method. Briefly, PEG-PLA-BODIPY were dissolved separately in diethylene glycol monoethyl ether at concentrations 25 mg/mL. Then, the polymer was mixed with 1 mg of Rhodamine B in a small glass vial. After this the solutions of polymers with dye were quickly nanoprecipitated in 5 mL of cold water and were kept for 5 min at 1000 r.p.m. stirring at room temperature. The NPs were then washed three times with cold water using Amicon® Ultra Centrifugal Filter (molecular weight cut-off - 3 kDa) to remove free Rhodamine B. FRET NPs were characterized by DLS, UV-vis, fluorescence and SEC HPLC. The FRET efficiency (E) was calculated using following equation: *E* = 1−*I*_DA_/*I*_D_. I_DA_ and I_D_ are the total donor fluorescence intensities in the presence and absence of the acceptor, respectively. The Forster distance (R_0_) was calculated using following equation: R_0_ = 0.002108(k^2^Φ_D_n^−4^J)^1/6^. The FRET orientation factor k^2^ = 2/3, the refractive index of PEG-PLA n = 1.46 the BODIPY-FL quantum yield Φ_D_ = 0.9, and the large spectral overlap integral J = 2.96 × 10^15^ nm^4^ M^−1^ cm^−1^.

### Intravital imaging

All animal procedures were approved by the Institutional Animal Care and Use Committee at the UMass Chan medical school at Worcester in accordance with all relevant guidelines. Bl6/C57 mice between the ages of 6 and 7 weeks were obtained from the Jackson Laboratory. Mice were anesthetized with 2% isoflurane and mounted on a Kopf stereotaxic frame. For open skull cranial window, the skull was drilled and thinned until the bone thickness can be easily pushed down and then gently removed by tweezers, leaving the dura intact. Then glass coverslip was placed over the surface of the craniotomy and fixed to the surface of the skull by gluing its edges to the head bone using cyanoacrylate glue. For thinned-skull cranial window, the skull was thinned carefully by bone files until it is transparent and thin enough to allow clear visualization of the underlying cortical vasculature without removing the bone entirely. Then glass coverslip was put on the cyanoacrylate glue over the window surface. To create an abdominal window, mice were anesthetized with 2% isoflurane, then a small incision was made through the skin and abdominal wall to expose the liver. After this thermo glass plate (Scintica) was used to optically expose and stabilize liver for imaging. To visualize the cerebral vasculature, 7 µL of Cy5 CD31 antibodies were injected in the tail vein. To inject NPs intravenously 50 µL of PEG-PLA-Cy5 NPs and PEG-PLA-BODIPY NPs separately were mixed with 100 µL of saline solution. For the injection of PEG-PLA-BODIPY NPs in the cisterna magna, 10 µL of NPs were prepared in aCSF (pH=7). The intravital images were collected using Scintica confocal microscope and exported to ImageJ for further analysis. To characterize the movement dynamics of nanoparticles, the mean square displacement (MSD) analysis was used. NPs were manually tracked 60 and 160 min postinjection by using ImageJ TrackMate software.

## Acknowledgments

We thank Dr. Gregory Stein and Curtana Pharmaceuticals for supplying CT179. This work was supported by funding from the National Institute for Neurological Disease and Stroke (NINDS) and the Eunice Kennedy Shriver National Instiute of Child Health and Disease (NICHD) at the National Institutes of Health (R01NS111292, R01HD099543, R01NS116657). Transmission Electron Microscopy (TEM) imaging was performed in the Electron Microscopy Core Facility at UMass Chan Medical School. We thank the Electron Microscopy Core Facility at UMass Chan Medical School, especially Gregory Hendricks, Ph.D. and Kyoung Hwan Lee for their assistance.

