## Supplemental Data for "Solvent-free Nanoparticle Assembly Protocol (SNAP): one-pot formulation of drug loaded polyester nanoparticles and their vessel size-dependent perivascular transport in the brain"

**Table S1.** Mean size, PDI, ζ-potential for NPs formed by PLA-PEG.

| **PEG-PLA**  **(5k-14k)** |  |  |  |
| --- | --- | --- | --- |
| **Conc., mg/ml** | **Size, mn** | **PDI** | **Ζ-potential, mV** |
| **25** | 79.24±2.32 | 0.124±0.016 | -13.60±2.09 |
| **50** | 96.90±8.96 | 0.197±0.016 | -14.88±1.56 |
| **100** | 106.35±3.15 | 0.176±0.013 | -16.64±2.05 |
| **PEG-PLA**  **(5k-18k)** |  |  |  |
| **Conc., mg/ml** | **Size, mn** | **PDI** | **Ζ-potential, mV** |
| **25** | 129.35±9.35 | 0.179±0.026 | -12.34±1.50 |
| **50** | 154.14±10.27 | 0.171±0.014 | -17.12±0.90 |
| **100** | 200.73±4.50 | 0.169±0.019 | -18.61±3.66 |
| **PEG-PLA**  **(5k-35k)** |  |  |  |
| **Conc., mg/ml** | **Size, mn** | **PDI** | **Ζ-potential, mV** |
| **25** | 137.81±14.11 | 0.147±0.025 | -13.49±1.64 |
| **50** | 141.68±3.34 | 0.141±0.017 | -18.82±1.41 |
| **100** | 204.46±7.18 | 0.159±0.019 | -20.73±3.30 |

**Table S2.** Mean size, PDI, ζ-potential for NPs formed by PCL-PEG.

| **PEG-PCL**  **(2k-5.2k)** |  |  |  |
| --- | --- | --- | --- |
| **Conc., mg/ml** | **Size, mn** | **PDI** | **Ζ-potential, mV** |
| **25** | 61.20±2.62 | 0.175±0.016 | -7.79±1.17 |
| **50** | 50.22±1.18 | 0.173±0.009 | -13.16±0.77 |
| **100** | 43.01±1.10 | 0.182±0.015 | -17.84±0.72 |
| **PEG-PCL**  **(5k-20k)** |  |  |  |
| **Conc., mg/ml** | **Size, mn** | **PDI** | **Ζ-potential, mV** |
| **25** | 142.96±16.81 | 0.180±0.020 | -6.80±0.32 |
| **50** | 133.66±3.74 | 0.154±0.016 | -10.97±0.53 |
| **100** | 151.09±3.47 | 0.127±0.017 | -14.41±0.39 |
| **PEG-PCL**  **(10k-40k)** |  |  |  |
| **Conc., mg/ml** | **Size, mn** | **PDI** | **Ζ-potential, mV** |
| **25** | 233.26±18.23 | 0.230±0.013 | -8.36±0.55 |
| **50** | 200.13±9.00 | 0.185±0.019 | -12.73±0.49 |
| **100** | 226.56±32.45 | 0.149±0.022 | -12.24±0.63 |

**Table S3.** Mean size, PDI, ζ-potential, loperamide loading and encapsulation efficiency for NPs formed by PLA-PEG.

| **PEG-PLA**  **(5k-14k)** |  |  |  |  |  |
| --- | --- | --- | --- | --- | --- |
| **Conc., mg/ml** | **Size, mn** | **PDI** | **Ζ-potential, mV** | **DL, %** | **EE, %** |
| **25** | 76.9±2.6 | 0.159±0.030 | -33.0±1.6 | 1.2±0.4 | 0.5±0.01 |
| **50** | 77.6±4.4 | 0.162±0.056 | -32.5±0.5 | 0.3±0.1 | 1.1±0.4 |
| **75** | 80.3±2.8 | 0.131±0.035 | -27.7±0.5 | 0.2±0.01 | 1.4±0.1 |
| **PEG-PLA-COOH (5k-16k)** |  |  |  |  |  |
| **Conc., mg/ml** | **Size, mn** | **PDI** | **Ζ-potential, mV** | **DL, %** | **EE, %** |
| **25** | 90.8±12.1 | 0.164±0.052 | -26.8±4.1 | 2.1±0.2 | 7.0±1.5 |
| **50** | 103.4±9.9 | 0.133±0.013 | -30.3±4.8 | 0.1±0.03 | 0.8±0.3 |
| **75** | 97.7±10.5 | 0.121±0.021 | -29.8±2.5 | 0.1±0.01 | 0.9±0.1 |
| **PEG-PLA**  **(5k-18k)** |  |  |  |  |  |
| **Conc., mg/ml** | **Size, mn** | **PDI** | **Ζ-potential, mV** | **DL, %** | **EE, %** |
| **25** | 97.7±11.6 | 0.194±0.09 | -34.9±2.2 | 0.2±0.01 | 0.4±0.1 |
| **50** | 106.8±2.8 | 0.104±0.019 | -31.9±0.7 | 0.1±0.008 | 0.8±0.2 |
| **75** | 115.4±4.8 | 0.143±0.016 | -32.4±2.8 | 0.2±0.001 | 1.3±0.2 |
| **PEG-PLA**  **(5k-35k)** |  |  |  |  |  |
| **Conc., mg/ml** | **Size, mn** | **PDI** | **Ζ-potential, mV** | **DL, %** | **EE, %** |
| **25** | 121.0±8.3 | 0.128±0.066 | -34.9±1.4 | 0.5±0.1 | 0.6±0.1 |
| **50** | 127.8±4.9 | 0.143±0.047 | -38.6±3.4 | 0.2±0.05 | 0.8±0.2 |
| **75** | 144.6±8.0 | 0.100±0.031 | -36.9±0.5 | 0.1±0.05 | 0.5±0.4 |

**Table S4.** Mean size, PDI, ζ-potential, loperamide loading and encapsulation efficiency for NPs formed by PCL-PEG.

| **PEG-PCL**  **(2k-5.2k)** |  |  |  |  |  |
| --- | --- | --- | --- | --- | --- |
| **Conc., mg/ml** | **Size, mn** | **PDI** | **Ζ-potential, mV** | **DL, %** | **EE, %** |
| **25** | 61.0±6.0 | 0.169±0.028 | -14.4±5.1 | 0.2±0.04 | 0.4±0.002 |
| **50** | 46.9±1.7 | 0.116±0.020 | -11.4±2.5 | 0.3±0.02 | 1.3±0.3 |
| **75** | 43.8±1.5 | 0.087±0.020 | -9.1±1.5 | 0.3±0.01 | 2.3±0.03 |
| **PEG-PCL**  **(5k-20k)** |  |  |  |  |  |
| **Conc., mg/ml** | **Size, mn** | **PDI** | **Ζ-potential, mV** | **DL, %** | **EE, %** |
| **25** | 86.5±4.8 | 0.182±0.052 | -17.8±3.8 | 0.7±0.2 | 1.0±0.3 |
| **50** | 137.6±41.7 | 0.185±0.051 | -25.3±2.3 | 0.4±0.07 | 2.2±0.8 |
| **75** | 109.3±3.9 | 0.127±0.021 | -26.3±0.7 | 0.4±0.01 | 3.3±0.4 |
| **PEG-PCL**  **(10k-40k)** |  |  |  |  |  |
| **Conc., mg/ml** | **Size, mn** | **PDI** | **Ζ-potential, mV** | **DL, %** | **EE, %** |
| **25** | 108.4±3.6 | 0.124±0.062 | -22.7±6.5 | 0.2±0.02 | 0.5±0.2 |
| **50** | 126.5±4.2 | 0.083±0.028 | -24.6±1.0 | 0.3±0.02 | 1.5±0.3 |
| **75** | 190.8±6.5 | 0.112±0.023 | -26.3±0.7 | 0.2±0.03 | 0.7±0.1 |

**Table S5.** Mean size, PDI, ζ-potential, loperamide and Rhodamine B loading and encapsulation efficiency for NPs formed by PLA-PEG.

| **PEG-PLA-BODIPY**  **(25mg/ml)** | **Size, nm** | **PDI** | **ζ-potential, mV** | **DL, %** | **EE, %** |
| --- | --- | --- | --- | --- | --- |
| **blank** | 79.4±16.8 | 0.159±0.047 | -30.4±3.1 | - | - |
| **loperamide** | 96.9±20.7 | 0.192±0.042 | -31.2±11.6 | 1.9±0.4 | 6.1±2.0 |
| **Rhodamine B** | 128.6±4.9 | 0.136±0.021 | -39.3±0.1 | 1.2±0.1 | 6.7±0.1 |

**Table S6.** Mean size, PDI, ζ-potential, drugs loading and encapsulation efficiency for NPs formed by PEG-PLA-COOH.

| **PEG-PLA-COOH (25mg/ml)** | **Size, nm** | **PDI** | **ζ-potential, mV** | **DL, %** | **EE, %** |
| --- | --- | --- | --- | --- | --- |
| **blank** | 87.4±5.8 | 0.146±0.019 | -25.4±6.7 | - | - |
| **diclofenac** | 106.3±10.1 | 0.256±0.022 | -37.7±6.5 | 5.1±0.8 | 12.9±5.8 |
| **loperamide** | 90.8±12.1 | 0.164±0.052 | -26.8±4.1 | 2.1±0.2 | 7.0±1.5 |
| **CT179** | 111.1±9.6 | 0.234±0.022 | -33.8±2.0 | 1.9±0.5 | 11.1±1.4 |
| **bortezomib** | 91.5±9.1 | 0.184±0.014 | -33.9±3.3 | 1.2±0.1 | 8.3±0.04 |
| **panobinostat** | 154.0±48.7 | 0.271±0.066 | -28.0±8.6 | 1.0±0.1 | 11.1±1.8 |
| **docetaxel** | 74.0±5.3 | 0.150±0.023 | -31.5±1.7 | 0.4±0.1 | 0.6±0.1 |
| **methotrexate** | 109.8±26.8 | 0.224±0.014 | -29.9±8.5 | 0.1±0.1 | 1.4±1.6 |
| **camptothecin** | 139.3±42.3 | 0.234±0.084 | -31.7±5.4 | 0.01±0.002 | 0 |

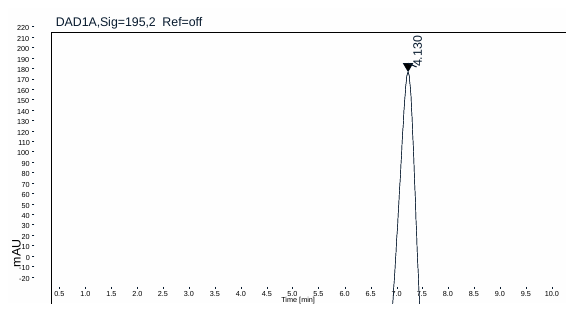

**Figure S1.** SEC-HPLC analysis for blank SNAP NPs.

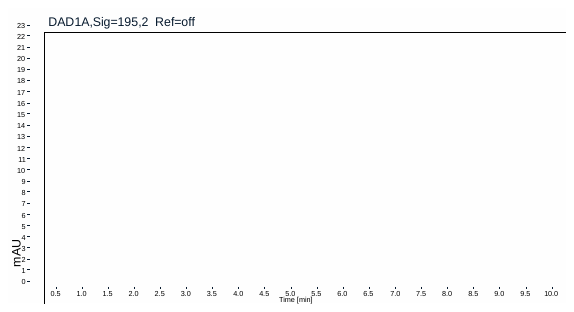

**Figure S2.** SEC-HPLC analysis for transcutol.

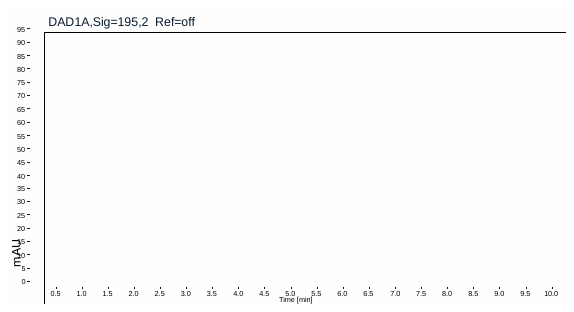

**Figure S3.** SEC-HPLC analysis for loperamide.

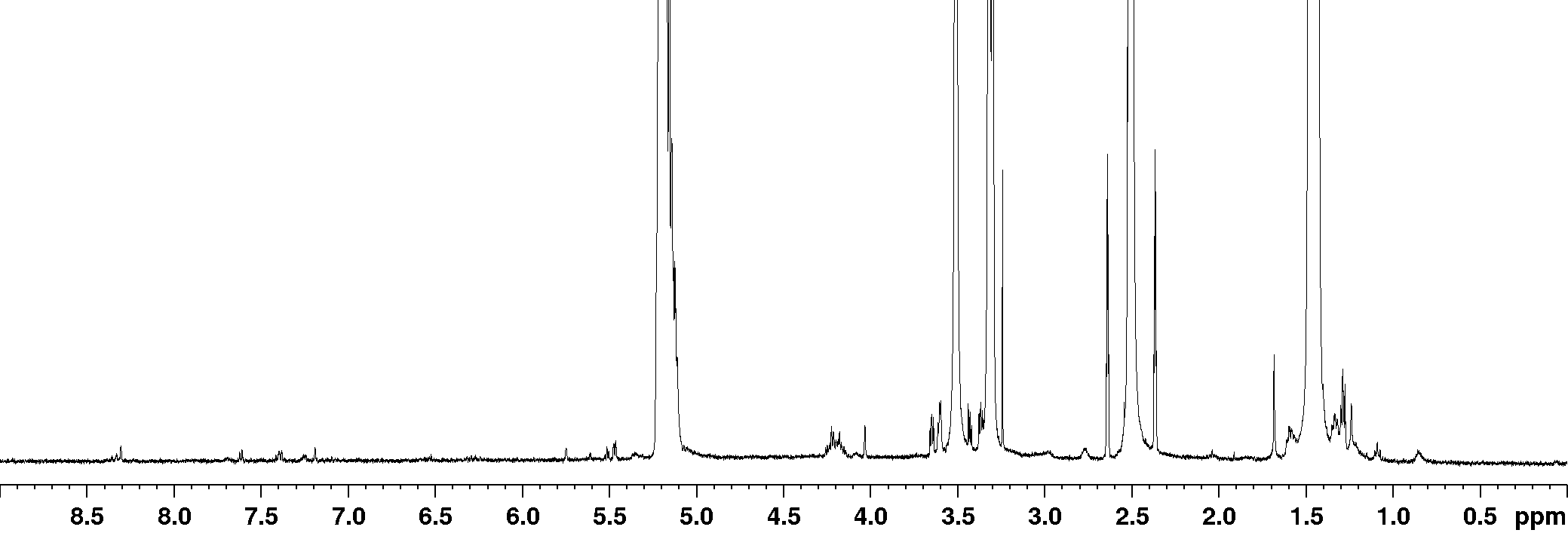

**Figure S4.** ^1^H-NMR spectrum in DMSO-d6 of Cy5-PLA-PEG.

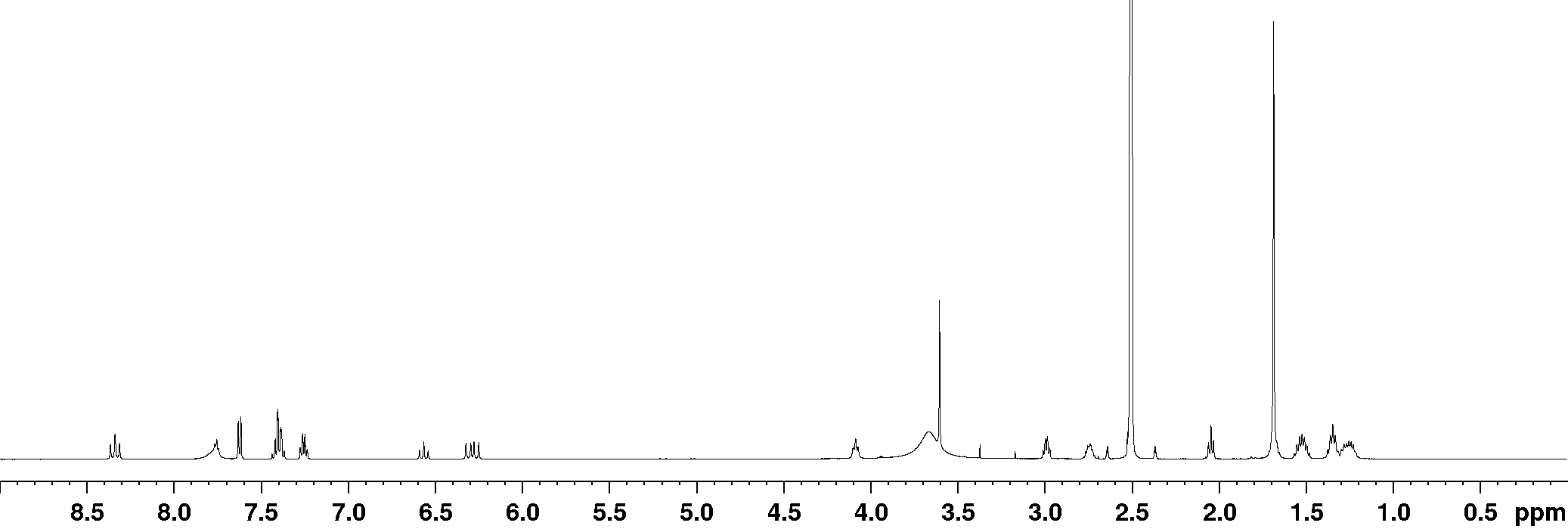

**Figure S5.** ^1^H-NMR spectrum in DMSO-d6 of Cy5-NH_2_

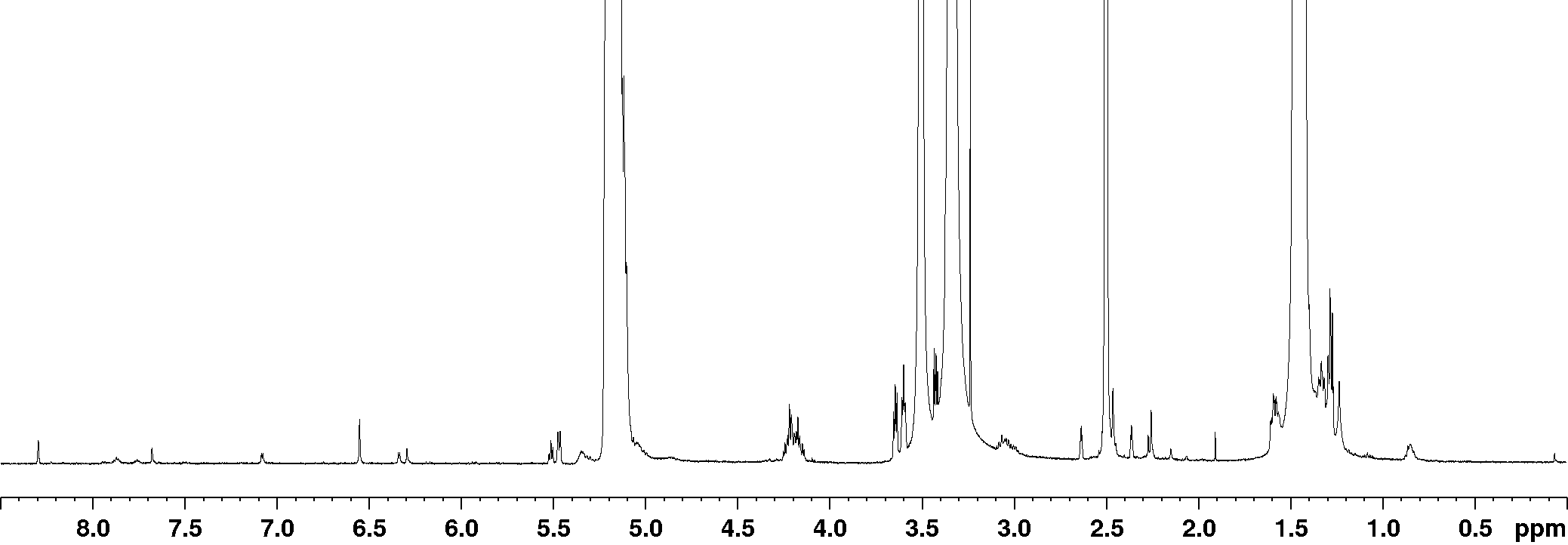

**Figure S6.** ^1^H-NMR spectrum in DMSO-d6 of PEG-PLA-BODIPY.

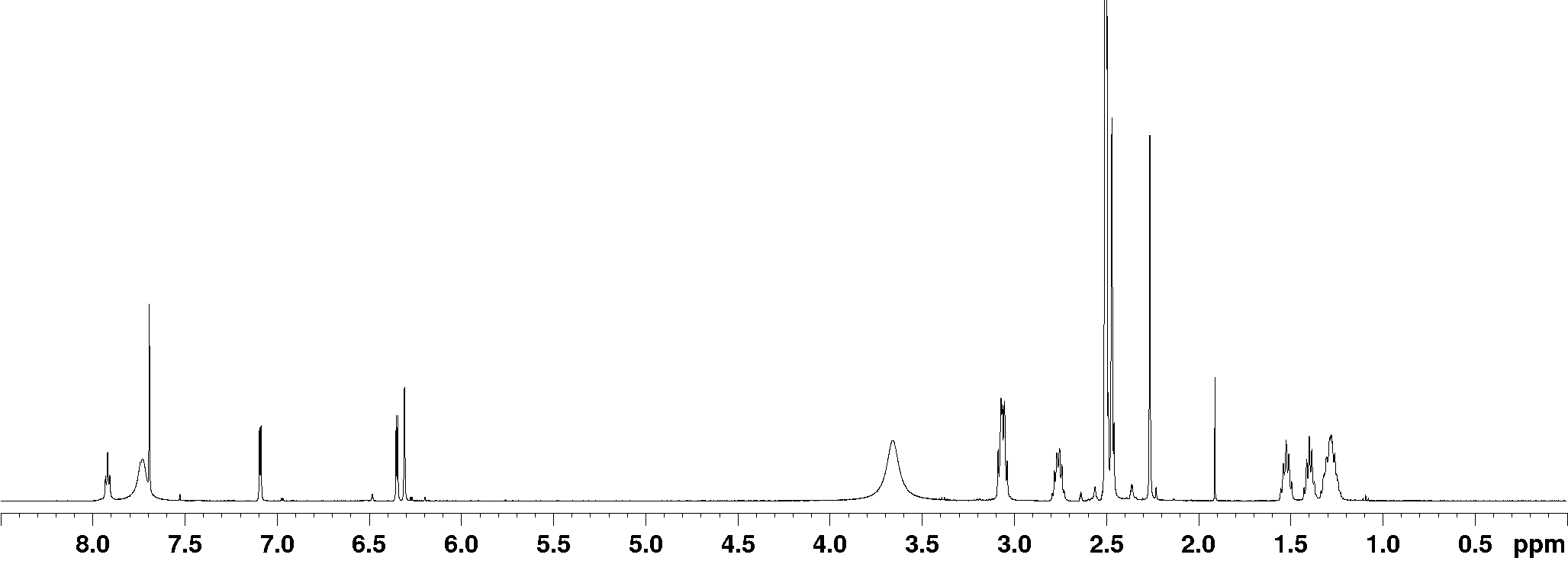

**Figure S7.** ^1^H-NMR spectrum in DMSO-d6 of BODIPY-FL-NH_2_

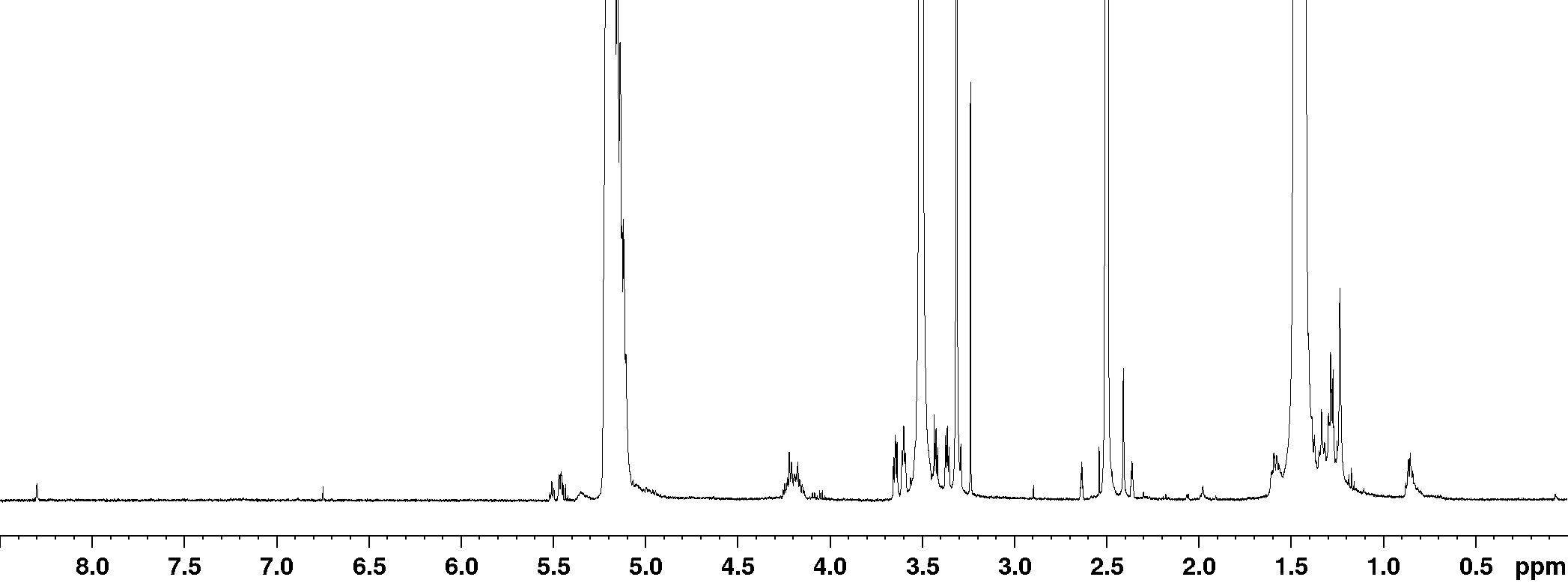

**Figure S8.** ^1^H-NMR spectrum in DMSO-d6 of PEG-PLA-COOH.

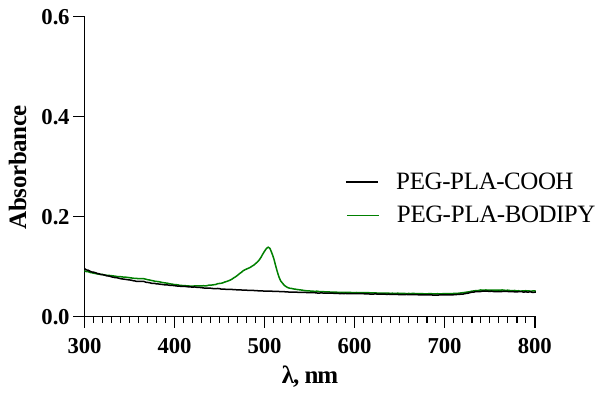

**Figure S9** UV-vis spectra for PEG-PLA-COOH and PEG-PLA-BODIPY NPs in water at 22˚C.

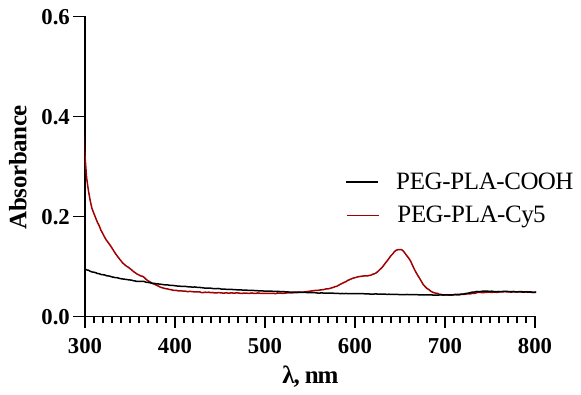

**Figure S10** UV-vis spectra for PEG-PLA-COOH and PEG-PLA-Cy5 NPs in water at 22˚C.

**Figure S11.** (A, B) Intravital imaging through an abdominal window of liver vasculature network labeled by CD31 (red) and distribution of NPs (green) based on PEG-PLA-C(O)NH-BODIPY delivered to the liver via intravenous injection into a tail vein. (C) The in vivo normalized median fluorescence intensity (nMFI) - post-injection time curve of NPs. (D, E) Intravital imaging through a thinned skull cranial window of brain vasculature network (medium brain vessels (mbv) and large brain vessels (lbv)) after NPs based on PEG-PLA-C(O)NH-Cy5 (green) administration into a tail vein. (F) The in vivo normalized median fluorescence intensity (nMFI) - post-injection time curve of NPs.
