## Supplementary figures and images for "Solvent-free Nanoparticle Assembly Protocol (SNAP): one-pot formulation of drug loaded polyester nanoparticles and their vessel size-dependent perivascular transport in the brain"

### Supplemental Data Movies

#### Slide 1
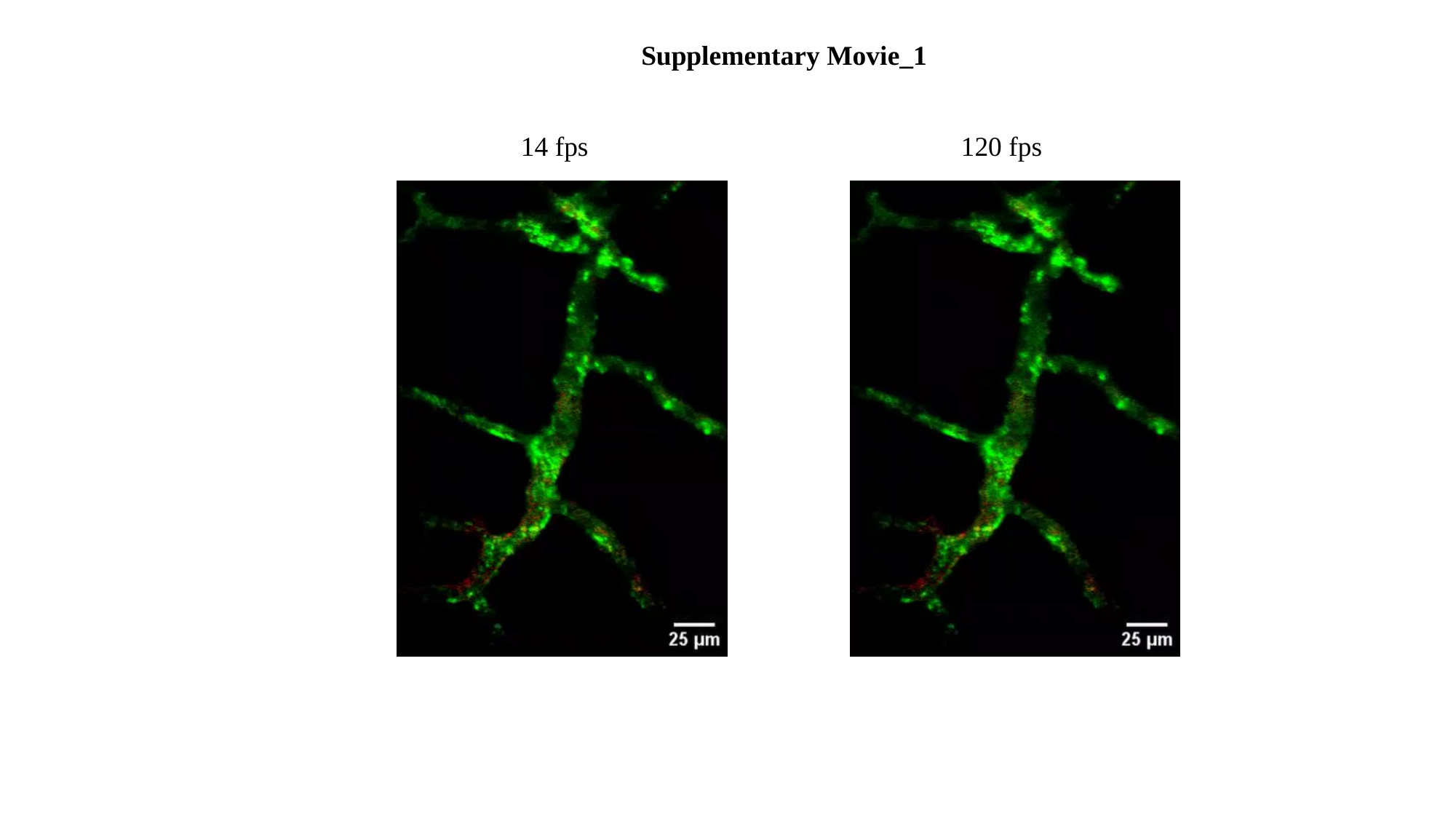

Supplementary Movie_1
14 fps 120 fps

#### Slide 2
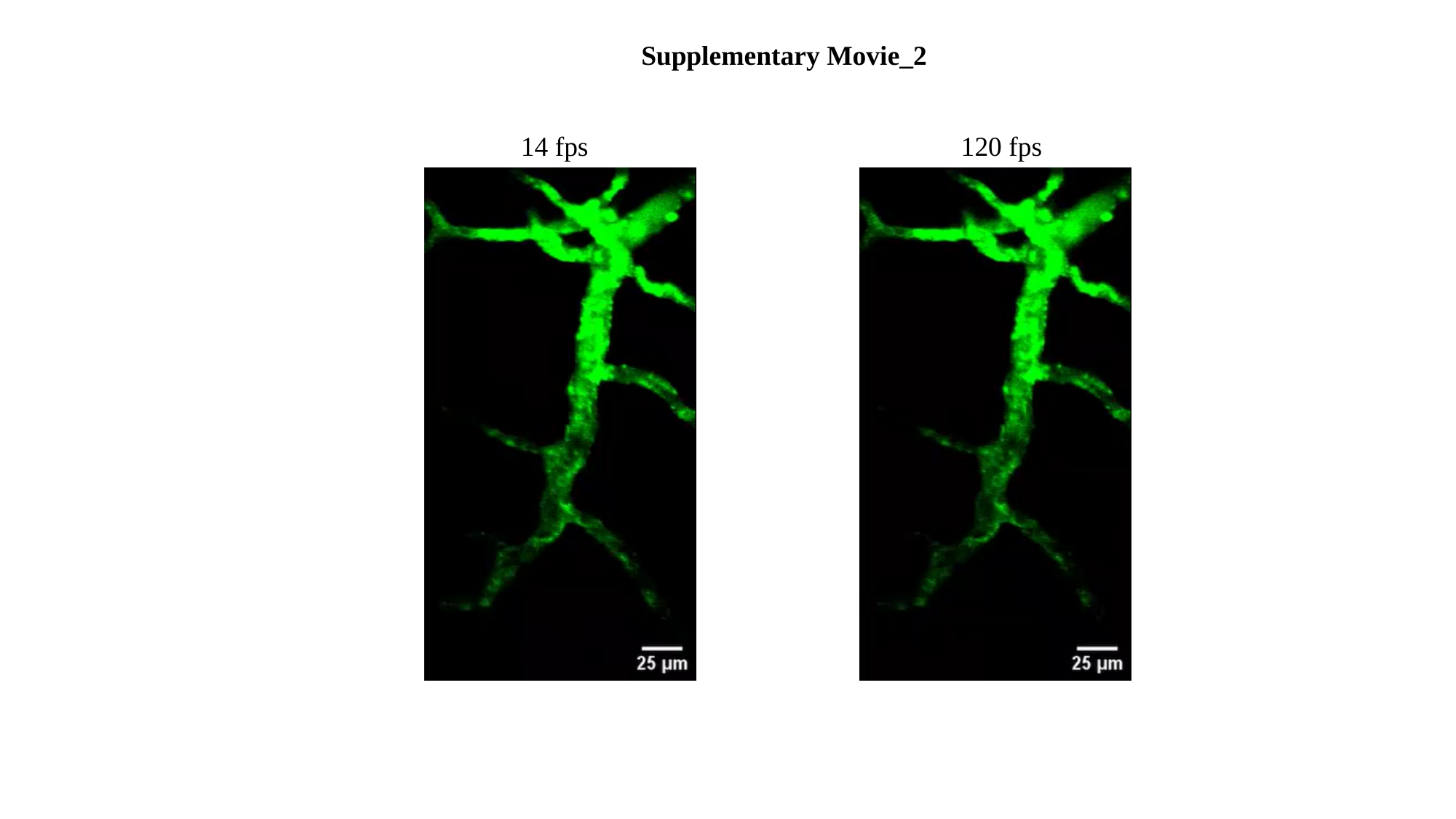

Supplementary Movie_2
14 fps 120 fps
